# Rapid and sensitive direct detection and identification of poliovirus from stool and environmental surveillance samples using nanopore sequencing

**DOI:** 10.1101/2020.04.27.053421

**Authors:** Alexander G. Shaw, Manasi Majumdar, Catherine Troman, Áine O’Toole, Blossom Benny, Dilip Abraham, Ira Praharaj, Gagandeep Kang, Salmaan Sharif, Muhammad Masroor Alam, Shahzad Shaukat, Mehar Angez, Adnan Khurshid, Nayab Mahmood, Yasir Arshad, Lubna Rehman, Ghulam Mujtaba, Ribqa Akthar, Muhammad Salman, Dimitra Klapsa, Yara Hajarha, Humayun Asghar, Ananda Bandyopadhyay, Andrew Rambaut, Javier Martin, Nicholas Grassly

**Affiliations:** Department of Infectious Disease Epidemiology, Imperial College London, London, W2 1PG, UK; Division of Virology, National Institute for Biological Standards and Control (NIBSC), South Mimms, Potters Bar, Herts EN6 3QG, UK; Institute of Evolutionary Biology, University of Edinburgh, Ashworth Laboratories, Edinburgh, EH9 3JT, UK; Division of Gastrointestinal Sciences, Christian Medical College, Vellore, India; Division of Epidemiology & Communicable Diseases, Indian Council of Medical Research, New Delhi, India; Department of Virology, National Institute for Health, Chak Shahzad, Islamabad, Pakistan; World Health Organization Eastern Mediterranean Regional Office, Amman, Jordan; Bill and Melinda Gates Foundation, 500 5th Ave N, Seattle, WA 98109, United States

## Abstract

Global poliovirus surveillance involves virus isolation from stool and environmental samples, intratypic differential (ITD) by PCR and sequencing of the VP1 region to distinguish vaccine (Sabin), vaccine-derived and wild-type polioviruses and ensure an appropriate response. This cell-culture algorithm takes 2-3 weeks on average between sample receipt and sequencing. Direct detection of viral RNA using PCR allows faster detection but has traditionally faced challenges related to poor sensitivity and difficulties in sequencing common samples containing poliovirus and enterovirus mixtures. We present a nested PCR and nanopore sequencing protocol that allows rapid (<3 days) and sensitive direct detection and sequencing of polioviruses in stool and environmental samples. We developed barcoded primers and a real-time analysis platform that generate accurate VP1 consensus sequences from multiplexed samples. The sensitivity and specificity compared with cell-culture were 90.9% (95% Confidence Interval: 75.7-98.1%) and 99.2% (95.5-100.0%) for wild-type 1 poliovirus, 92.5% (79.6-98.4%) and 98.7% (95.4-99.8%) for vaccine and vaccine-derived serotype 2 poliovirus, and 88.3% (81.2-93.5%) and 93.2% (88.6-96.3%) for Sabin 1 and 3 poliovirus alone or in mixtures when tested on 155 stool samples in Pakistan. Variant analysis of sequencing reads also allowed identification of polioviruses and enteroviruses in artificial mixtures and was able to distinguish complex mixtures of polioviruses in environmental samples. The median identity of consensus nanopore sequences with Sanger or Illumina sequences from the same samples was >99.9%. This novel method shows promise as a faster and safer alternative to cell-culture for the detection and real-time sequencing of polioviruses in stool and environmental samples.

## Introduction

Resurgent wild-type poliovirus transmission in Pakistan and Afghanistan, and expanding outbreaks of vaccine-derived poliovirus (VDPV) in Africa pose the risk of reversing progress made by the Global Polio Eradication Initiative. Effective control of these outbreaks using oral poliovirus vaccine relies on rapid detection in stool samples from children with acute flaccid paralysis (AFP) or from environmental surveillance (ES) samples. Detection of poliovirus is not confirmed until a genetic sequence from the VP1 region is obtained, which allows circulating virus to be distinguished from vaccine strains.

The current gold standard method of poliovirus detection is by culture of the virus using susceptible cell lines, reverse transcription PCR (RT-PCR) of RNA extracted from cell-culture supernatant to determine serotype and distinguish Sabin from non-Sabin polioviruses (intratypic differentiation (ITD)), and subsequent sequencing by traditional (Sanger) methods (1). However, this method is slow and leads to delays between sample collection and sequencing result, compromising the speed and effectiveness of any vaccination response. For example, during 2016-18 the median time taken to obtain a sequence needed for case confirmation varied from three to six weeks in countries in Africa and Asia with VDPV or wild-type poliovirus outbreaks (2). These delays reflect both the complex and time-consuming cell-culture steps, but also in many countries the need to ship samples internationally to laboratories with sequencing facilities. These methods have other limitations, including challenges related to biocontainment of poliovirus grown in culture, virus mutation during growth in cell-culture and competitive exclusion of viruses in samples containing poliovirus mixtures leading to a lack of detection of minority or low fitness variants (e.g. ES samples containing vaccine (Sabin) and vaccine-derived polioviruses).

Direct molecular detection of polioviruses in stool or ES samples using RT-PCR could allow rapid detection of poliovirus without the need for cell-culture. However, quantitative RT-PCR capable of amplifying all polioviruses through use of degenerate (pan-poliovirus) primers has historically reported low sensitivity compared with cell-culture (3). Use of serotype or strain-specific primers improves sensitivity, but multiplex assays with many primers would be required to identify all polioviruses and the resulting products are too short for informative sequencing (4). Quantitative PCR in the absence of confirmatory sequencing also risks false positive results because of cross-contamination, or non-specific amplification when using degenerate primers (5).

A sensitive, nested RT-PCR for direct detection of polioviruses and other enteroviruses from clinical material has been developed and recommended by the World Health Organisation (WHO) for enterovirus surveillance, but this method generates only partial VP1 sequences that cannot be used to confirm poliovirus infection (6). More recently, a nested RT-PCR based on amplification of the entire capsid region using pan-enterovirus primers, followed by the full VP1 using degenerate pan-poliovirus primers was shown to be highly sensitive for detection of poliovirus in stool (3). However, sequencing of the VP1 product using the traditional (Sanger) method is problematic for samples containing >1 poliovirus because of its inability to accurately sequence mixed samples. Similarly, non-specific amplification of VP1 from other enteroviruses, which are highly prevalent in stool in low-income countries and practically ubiquitous in ES samples, limits the applicability of this approach.

Here we present a method for direct detection and identification of polioviruses from stool and ES samples using a nested PCR followed by nanopore (MinION) sequencing of the poliovirus VP1 region. This approach allows rapid sequencing library preparation following RNA extraction (<14 hours) and subsequent, simultaneous sequencing of multiple samples using a barcoded DNA library. Sequences are demultiplexed and analysed in real-time to provide immediate identification of poliovirus. We validated the method using artificial mixtures of enteroviruses and assessed sensitivity and specificity against culture and quantitative PCR using stool and ES samples from Pakistan and India.

## Materials and Methods

### Creation of artificial virus mixtures

We used virus stocks of Echovirus 7 (E7), Enterovirus-A71 (A71), Enterovirus-D68 (D68) and serotype 1 Sabin poliovirus (PV1) containing 4.10 × 10^9^, 4.39 × 10^8^, 1.26 × 10^7^ and 4.00 × 10^9^ median cell-culture infectious doses (CCID_50_)/ml, respectively. We prepared an even mixture by adding 20 ul cell-culture supernatant of E7, 200 ul of EV71, 1000 ul of D68 and 20 ul of PV1. To create an uneven sample, we mixed 20 ul of each and made the volume to 500 ul. Sample 3 was a pure culture of PV1 and Sample 4 was a trivalent oral poliovirus vaccine standard (7). We also tested novel oral poliovirus vaccine candidates that have been engineered for greater stability compared with Sabin vaccine (2 candidates for each serotype), including novel serotype 2 vaccine (nOPV2) that is scheduled for use in response to serotype 2 VDPV outbreaks under WHO Emergency Use Listing in 2020 (8, 9).

### Stool and environmental samples

Stool samples were collected during acute flaccid paralysis (AFP) surveillance in Pakistan between October 2019 and February 2020. We tested 155 samples found to be positive during this period by standard cell-culture and ITD for wild-type 1 poliovirus (33), circulating serotype 2 VDPV (11) or Sabin poliovirus serotype 1, 2 and/or 3 (111).

We additionally tested stored faecal aliquots from 1-4 year old Indian infants participating in a clinical trial of inactivated poliovirus vaccine conducted in 2013 and who had received bivalent OPV containing serotypes 1 and 3 poliovirus (10). In 2013/14 these samples were tested using a singleplex quantitative real-time reverse-transcription PCR (qRT-PCR) for Sabin poliovirus 1 and 3 (11) and a subset by cell-culture following the standard WHO protocol (12). We retrieved all 217 samples from this subset that were positive for serotype 1 or 3 poliovirus by qPCR and 49 randomly selected samples that were negative by qPCR and culture for inclusion in the current study.

Sewage samples collected by the grab method in 2015 and 2017 in Pakistan were also used. One-litre raw sewage specimens were processed using two-phase separation to generate sewage concentrate, followed by the poliovirus isolation cell culture algorithm following WHO guidelines (13). Results from six culture flasks were amalgamated using aliquots drawn from a single sample (3ml in total). Sewage concentrates were also used for direct PCR and sequencing using nanopore, Illumina or Sanger methods.

Institutional ethics approval was not sought because samples are anonymized and free of personally identifiable information. Samples from Pakistan were collected as part of the WHO/Pakistan Government’s official polio surveillance process.

### RNA extraction from stool and environmental sample concentrates

Viral RNA extracted from stool and environmental samples needed to yield long PCR products (~ 4000 bp). Based on previous experience with different commercially available RNA extraction kits (see Appendix S1), we established that both Roche and Qiagen kits were suitable (see Appendix S1). Roche Kit (Cat no. 11858882001) was used for RNA extractions from artificial mixtures and sewage samples (using 200 ul of sewage concentrate as input). For stool samples, 1g of faeces was added to 5 ml of PBS. Chloroform extractions were then performed as described in (14) and 140 ul of suspension was taken forward for RNA extraction using the QIAamp viral RNA mini kit in Pakistan and QIAamp 96 Virus QIAcube HT Kit in India.

### PCR amplification

PCR products for Sanger, Illumina and nanopore (MinION) sequencing were obtained using a combination of protocols. RT-PCR was performed using the Superscript III One-Step RT-PCR System (Invitrogen). cDNA was first synthesised with a single primer being added and the second primer added for the PCR step. RT-PCR products spanning the enterovirus capsid region or whole poliovirus genome were generated using primer combinations shown in Dataset S1 and Figure S1. Poliovirus VP1 products were generated by nested PCR, first amplifying the capsid region by RT-PCR with pan-enterovirus (pan-EV) primers followed by a second PCR using pan-poliovirus (pan-PV) primers Q8/Y7 or Q8/Y7R, or serotype-specific VP1 primers (Dataset S1 and Figure S1). DreamTaq DNA Polymerase (Thermo Scientific) or LongAmp Taq Polymerase (NEB) was used for the second PCR reaction.

### Sequencing library preparation for nanopore sequencing

As initial tests demonstrated strong amplification of VP1 using the pan-PV Q8/Y7 primers, we sought to reduce the time and cost for nanopore sequencing library preparation by modifying these primers with either the addition of a barcode adaptor (VP1-BCA) or barcode (VP1-barcode) to allow immediate multiplexing of samples (Figure 1). PCR products underwent standard preparation for nanopore sequencing with the LSK-109 kit with barcoding using the EXP-PBC096 kit where necessary. Products amplified with adaptor primers entered the library preparation protocol at the clean-up step after adaptor ligation. Products amplified with the barcoded primers entered the protocol at the clean-up step after the barcoding PCR. A complete protocol is documented in Appendix S2. Sequencing was performed on Oxford Nanopore Technologies (ONT) MinION Mk1B sequencers using R9.4 flow cells. Sequencing run conditions and outputs are described in Table S1.

**Figure 1.**
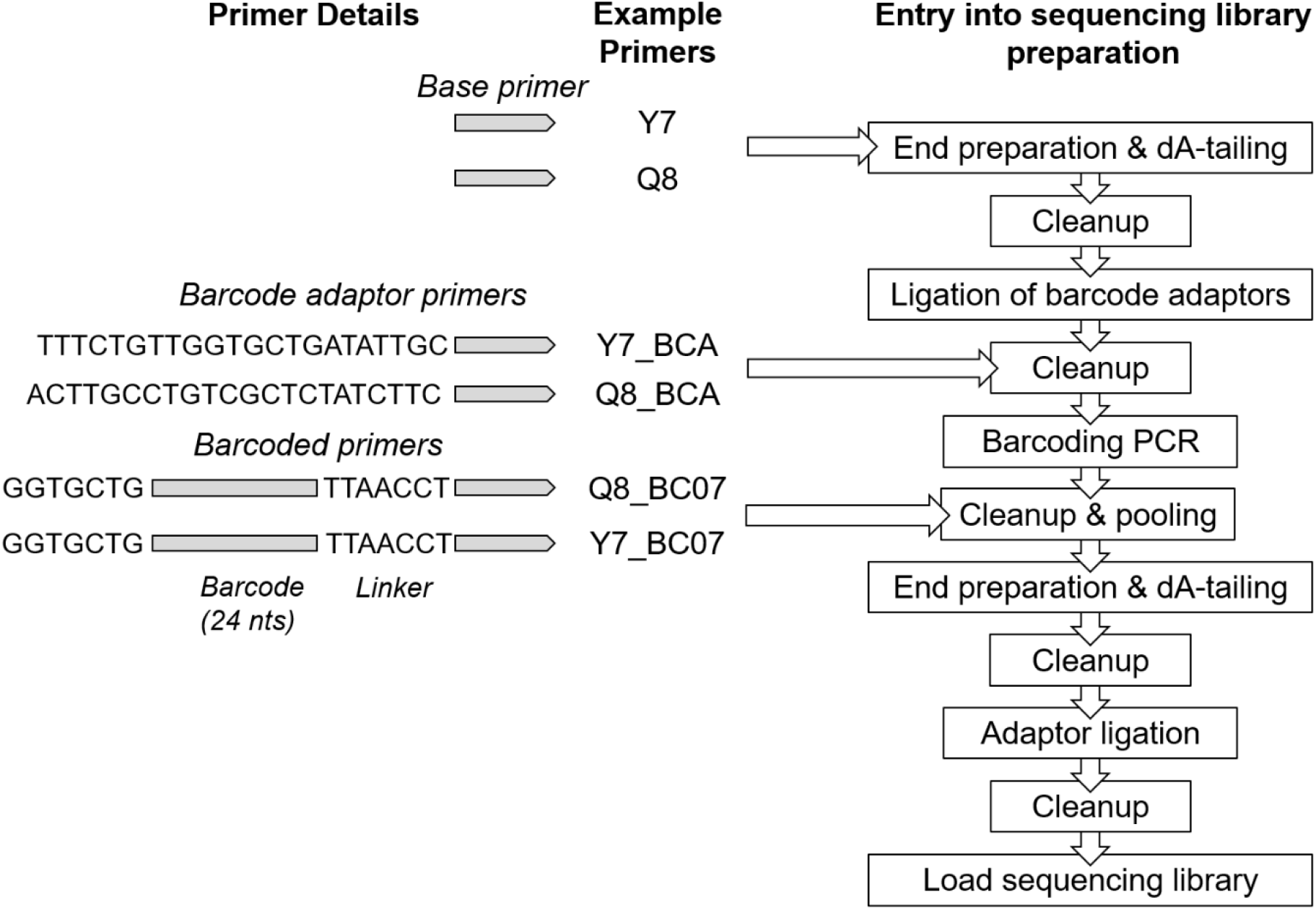
The formulation of primers for MinION sequencing and the entry of the various PCR products into the sequencing library preparation protocol. In addition to the basic primers, we have tested the addition of barcode adaptor (BCA primers) or barcodes and flanking regions (BC# primers, with # being the barcode number with reference to nanopore PCR ligation barcode sequences).

Pan-EV capsid and pan-PV whole-genome PCR products from the artificial mixtures and ES samples underwent the standard preparation for nanopore sequencing with the LSK-108 kit with barcoding using the EXP-PBC001 kit.

### Sanger sequencing of poliovirus VP1 nested PCR products

Amplified poliovirus VP1 products from nested PCR were purified using QIAquick PCR purification kit (Qiagen) and sequenced using an ABI Prism 3130 genetic analyser (Applied Biosystems) using standard protocols.

### Sequence library preparation for Illumina sequencing

Pan-EV capsid and pan-PV VP1 nested PCR products from ES samples were additionally sequenced using Illumina next-generation sequencing. Sequencing libraries were prepared using Nextera XT reagents and sequenced on a MiSeq using a 2×250-mer paired-end v2 flow cell and manufacturer’s protocols (Illumina).

### Bioinformatics

Nanopore data were drawn from twenty individual MinION sequencing runs, each of which is documented in Table S1. Fast5 files were base-called using Albacore v1.8 (pan-PV and pan-EV experiments) or Guppy version v3.4 (remaining experiments) with GPU acceleration on a Precision 7540 Dell laptop running Ubuntu 18 LTS with 32GB RAM, a 16 core Intel Xeon CPU and Nvidia RTX3000 GPU. Real-time analysis of VP1 reads was performed using a pipeline customised for the identification and mapping of PV nanopore data and then visualised using RAMPART (15). Once sufficient coverage of the VP1 region for each demultiplexed sample was confirmed using RAMPART, consensus sequences were generated from the base-called reads using a custom pipeline build on a Snakemake framework (16). In brief, reads are binned by sample and matched against a custom database comprising poliovirus and non-polio enterovirus (NPEV) VP1 regions drawn from NCBI Nucleotide (17) and the NIAID Virus Pathogen Database and Analysis Resource (ViPR) (18). For scoring of poliovirus presence, a minimum of 50 reads was required within a bin which is above the minimum threshold of sequencing reads required to give an accurate consensus as established with simulated data (see Figure S2). A read-correction (‘polishing’) module, which uses mafft v7 (19), minimap2 v 2.17 (20), racon v1.4.7 (21), medaka v0.10.0 (22), and python code for quality control and curation, creates a consensus sequence for each virus population. The installable, documented pipeline can be found at https://github.com/polio-nanopore/realtime-polio.

VDPV and other Sabin-related poliovirus sequences were not included in the VP1 reference database used by RAMPART because of the potential for spurious matches by nanopore-sequenced Sabin poliovirus given the sequencing error rate before polishing and consensus sequence generation. Instead, sequencing reads with a closest match to the reference Sabin poliovirus VP1 sequences for each serotype were classified as ‘Sabin-related’ polioviruses. Consensus sequences were subsequently generated for these Sabin-related reads to distinguish VDPV from Sabin polioviruses.

We additionally developed a bioinformatics pipeline to distinguish mixtures of polioviruses including VDPV and Sabin polioviruses of the same serotype. Raw reads were clustered at 85 % identity with vsearch v2.7.0 (23) and aligned to Sabin references using minimap2 v2.17. Variant calling was performed within clusters using medaka v0.10.0. Raw reads were then mapped to the variant sites to confirm linkage and a maximum likelihood tree of the representative variants generated using the phangorn package in R v3.4.3 (24, 25). Trees were cut at a height of 0.1, a consensus sequence for each subtree was generated and any identical consensus sequences were merged. Further development of this pipeline is ongoing and documented at https://github.com/polio-nanopore/realtime-polio. This same pipeline was used to detect potential cross-contamination of samples during RNA extraction or PCR that could result in a mixture of poliovirus reads for a single reaction.

Nanopore sequences from pan-EV capsid and pan-PV complete genome amplicons generated from artificial mixtures and ES samples were demultiplexed and trimmed using Porechop v0.2.3 and reads containing internal adaptors discarded. Reads were screened to retain only those of the expected length (3,500 to 4,200 nucleotides for the capsid and 7,000 to 7,700 nucleotides for the nearly full-length genome). Reads were mapped with blastn (26) to the custom VP1 database.

Filtered reads from Illumina sequencing of pan-EV and nested VP1 products generated with pan-PV Q8/Y7 primers were mapped to the VP1 database and PV consensus sequences extracted. These sequences data were processed and analysed using a custom workflow with Geneious R10 software package (Biomatters) as described before (27).

Phylogenetic trees showing the relatedness of VP1 sequences generated using different sequencing approaches were constructed in MEGAX (28) using the neighbour-joining method from evolutionary distances estimated under the Tamura-Nei substitution model using the maximum composite likelihood (29).

Demultiplexed sequencing data is available at the European Nucleotide Archive under accession number PRJEB37193.

## Results

### Identification of enteroviruses and polioviruses in artificial mixtures and ES samples using one-step RT-PCR and nanopore sequencing

Nanopore sequencing of the enterovirus capsid generated by RT-PCR using pan-EV primers identified all viruses included in the artificial mixtures, with read counts broadly reflecting the known composition of the mixtures (Figure 2a). Sabin poliovirus 1 was detected even when the virus constituted only 0.1% of the viral mixture. Discrepancy between the proportion of reads by serotype and known mixture of the even sample may derive from the pooling of products from the two pan-EV primer sets, with over-representation of species B and D enteroviruses (E7 and EV-D68). Nanopore sequencing of the pan-PV full genome product generated by RT-PCR of artificial poliovirus mixtures generated sequences for nearly the entire genome, with the abundance of reads closely matching that expected (Figure 2b).

**Figure 2.**
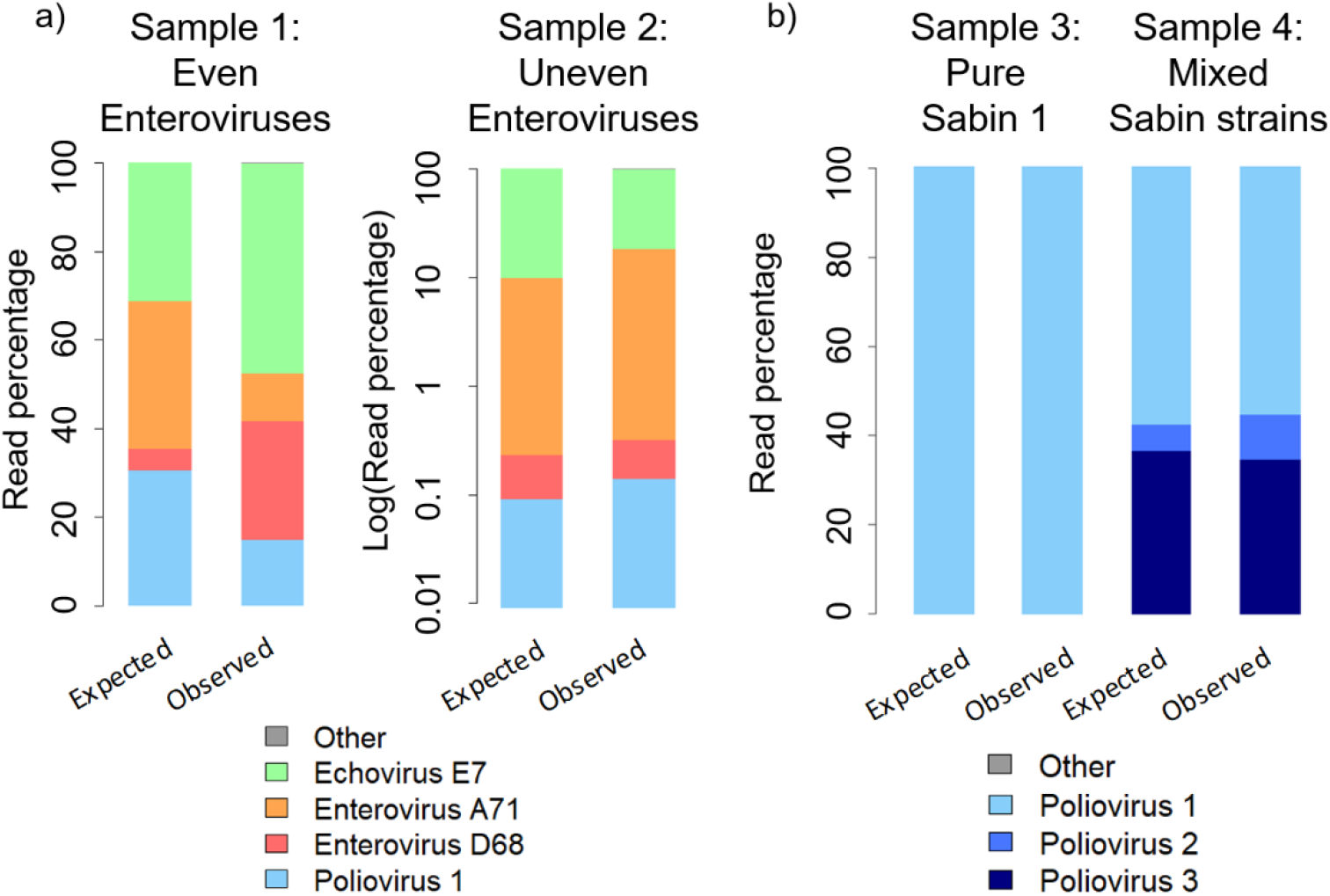
Identification of enterovirus and poliovirus mixtures by nanopore sequencing of the entire capsid or full genome. The observed abundance of sequencing reads is compared with that expected based on the known composition of a) artificial enteroviruses mixtures and b) poliovirus mixtures. For each sample the left bar shows the expected outcome according to the viral titres and the right bar shows the observed results through comparison of the sequencing reads with the VP1 database. The results for sample 2 have been plotted on a log scale.

We sequenced pan-EV RT-PCR entire capsid products from eight ES samples collected in Pakistan during 2015 and 2017 using nanopore and compared with assembled reads for the same region generated though Illumina sequencing. The distribution of enterovirus serotypes was highly similar for the two sequencing methods (Figure 3). Poliovirus was found in all eight samples by nanopore sequencing, yet attributed reads formed only a small percentage of total reads (0.002 to 6.65 %, median 0.03%). Whole-capsid poliovirus contigs were only produced for two of the eight ES samples from the Illumina reads, representing 0.08% and 5.47% of total reads, respectively.

**Figure 3.**
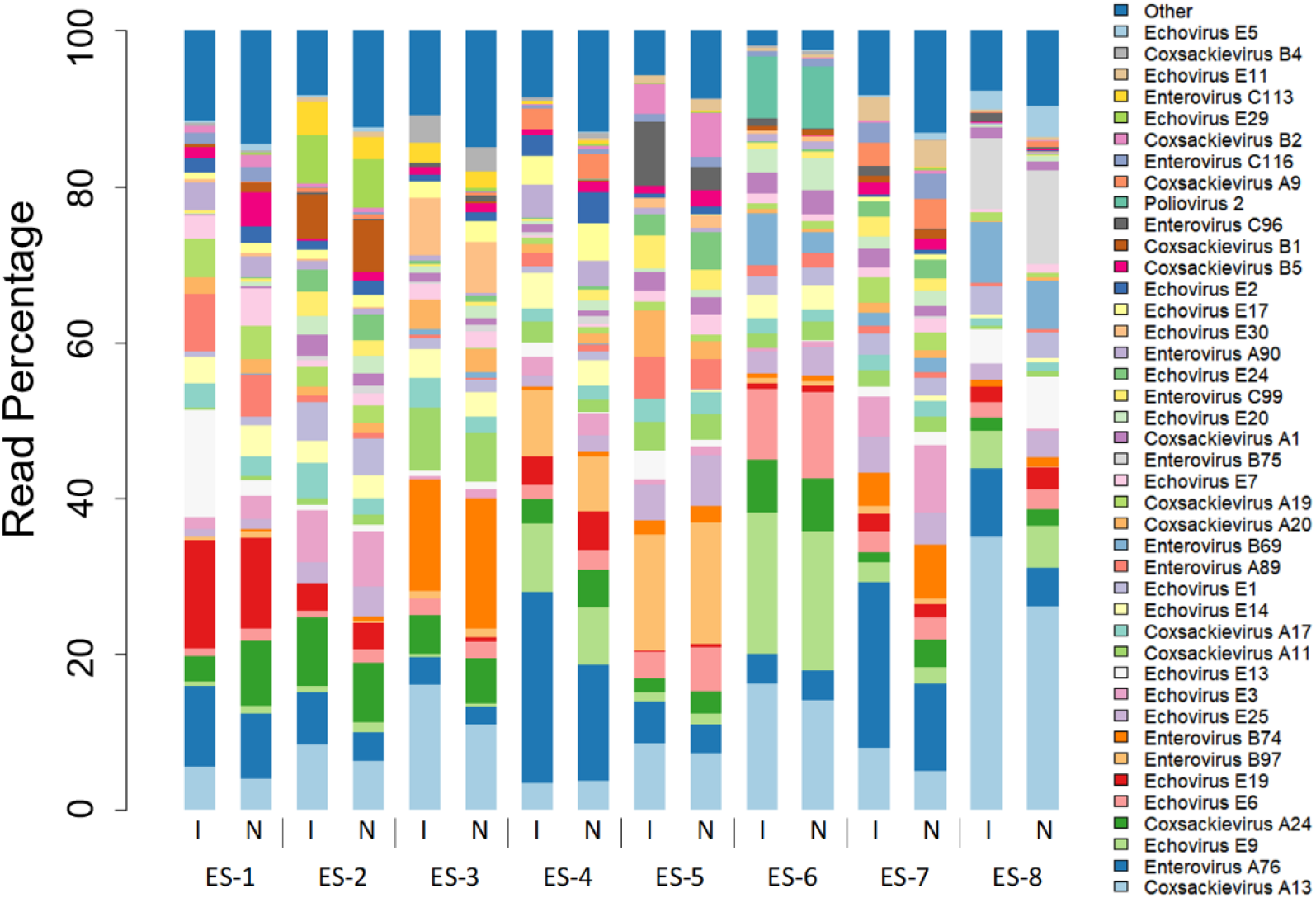
Enteroviruses present in eight ES samples from Pakistan analysed by nanopore (N) and Illumina Miseq (I) sequencing of pan-EV PCR products. Sequencing reads attributed to the 90% most abundant viruses are shown, with the remaining identified reads being grouped into “Other”.

### Nested capsid/VP1 PCR barcoded primer development

To increase the specificity and sensitivity of our method we performed a nested PCR, using pan-EV entire capsid RT-PCR with Cre-R and 5′NCR primers in an RT-PCR, followed by pan-PV PCR using Q8/Y7 VP1 primers (Dataset S1). Modification of the VP1 primers with the addition of ONT barcodes or the barcode adaptor sequence gave a slight decrease in sensitivity, but this was equivalent to the loss of product that occurred during the additional library preparation steps when using the unadapted primers (Appendix S3). Equivalence of the VP1-BCA and VP1-barcode primers was further confirmed by amplification of VP1 from the pan-EV products of 20 ES samples, which yielded DNA at >25 ng/µL suitable for sequencing for 18/20 (90%) and 20/20 (100%) of samples for each primer set respectively (Fisher’s exact test, p-value=0.487).

### Nested PCR and nanopore sequencing of stool samples

Nested PCR with barcoded Q8/Y7 primers and nanopore sequencing of the 155 stool samples collected during 2019-2020 in Pakistan detected 90.9% (30/33) of the wild-type 1 polioviruses that were identified by cell-culture and Sanger sequencing, 92.5% (37/40) of serotype 2 Sabin and VDPVs and 88.3% (106/120) of Sabin 1 and 3 polioviruses (Table 1). One wild-type poliovirus and one serotype 2 VDPV were detected in samples that were not positive for these viruses by culture and Sanger sequencing (giving specificity compared with culture of 99.2% and 99.3 % for these virus types respectively). Serotype 1, 2 and 3 Sabin polioviruses were detected in 14 samples that were not positive for these viruses by cell-culture (specificity 95.6%). The variant calling pipeline identified contamination in 16 samples (14 in the last 2 sequencing runs) and these reads were removed. For the 30 samples where wild-type 1 poliovirus was identified by both culture and nanopore sequencing the median identity for the VP1 region was 99.9 % (range: 99.0 % - 100 %; Figure 4).

**Table 1.** Detection and sequencing of poliovirus in 155 Pakistan stool samples using nanopore compared with cell culture and sequencing results

| **Culture Results** | **Nanopore sequencing results** |  |  |  |  |  |  |  |  |
| --- | --- | --- | --- | --- | --- | --- | --- | --- | --- |
|  | Sabin 1 | Sabin 2 | VDPV2 | Sabin 3 | Wild-type 1 | Sabin 1+3 | Sabin 1+3 + VDPV2 | Wild-type 1 + Sabin1 | Negative |
| Sabin 1 | **6** | 0 | 0 | 0 | 0 | 4 | 0 | 0 | 1 |
| Sabin 2 | 0 | **28** | 0 | 0 | 0 | 0 | 0 | 0 | 1 |
| VDPV2 | 0 | 1 | **9** | 0 | 0 | 0 | 0 | 0 | 1 |
| Sabin 3 | 0 | 0 | 0 | **24** | 1 | 5 | 1 | 0 | 2 |
| Wild-type 1 | 0 | 0 | 0 | 0 | **29** | 1 | 0 | 1 | 2 |
| Sabin 1+3 | 0 | 0 | 0 | 2 | 0 | **32** | 0 | 0 | 4 |
| Negative | 0 | 0 | 0 | 0 | 0 | 0 | 0 | 0 | 0 |

**Figure 4.**
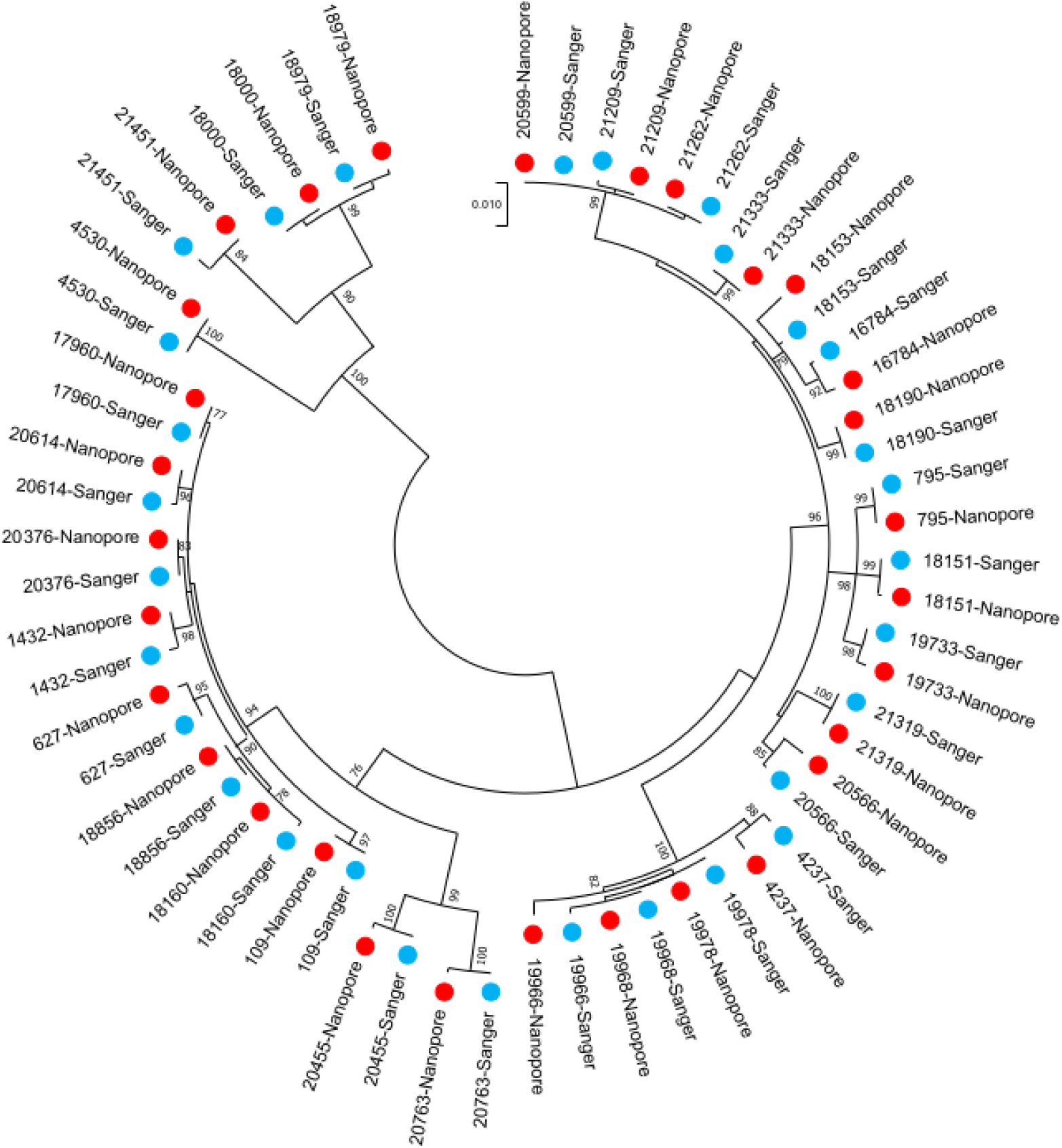
Relatedness of wild-type polioviruses 1 identified in stool samples in Pakistan using alternative methods. VP1 sequences based on direct PCR and nanopore sequencing (red) are compared with Sanger sequenced cell culture isolates (blue).

Stool samples containing serotype 1 or 3 Sabin poliovirus in India likely had suffered degradation of viral RNA as a result of storage for six years. Nonetheless, 69% (149/217) yielded >50 reads of the correct virus and this increased to 91% (109/120) among those with at least 100 copies of the target sequence at the time of original quantitative PCR testing (Appendix S4).

### Nested PCR and nanopore sequencing of novel oral poliovirus vaccine candidates

Nested PCR using Q8/Y7 primers generated a VP1 product for all eight novel oral poliovirus vaccine candidates and consensus sequences generated from nanopore reads were 100% identical to Illumina sequences generated for the same samples.

### Nested PCR and nanopore sequencing of ES samples

Nanopore sequencing using nested PCR with pan-PV VP1 Q8/Y7 primers resulted in a median of 87.1 % of VP1 enterovirus reads mapping to poliovirus (range 27.4% to 100.0%). This compared with just 6.8 % (range 0.1% to 95.9%) when using the more degenerate Q8/Y7R primer. Consensus building and variant calling and results from the more specific barcoded Q8/Y7 primers allowed the identification of poliovirus mixtures in these samples including wild-type, vaccine (Sabin) and vaccine-derived polioviruses corresponding to different serotypes (Figure 5). These results were almost identical to those obtained by Illumina sequencing of VP1 (Q8/Y7) nested PCR products and Sanger sequencing of VP1 nested PCR products generated with serotype-specific primers (Figure 4). Nanopore sequencing detected eight of nine wild-type 1 polioviruses identified by Sanger or Illumina sequencing of direct VP1 products and showed very similar detection of Sabin-related polioviruses (Pearson’s phi coefficients = 0.89, 1.0 and 1.0 for serotypes 1, 2 and 3 respectively for nanopore vs Sanger or Illumina results (which were identical)). In addition, it was possible to distinguish serotype 2 VDPV and Sabin polioviruses from nanopore sequencing data using our bioinformatics pipeline, which is not possible for Sanger or Illumina reads (the latter because they are short and it is difficult to confirm mutations are shared by a single virus). All three direct detection and sequencing methods gave somewhat different results from those generated by Sanger sequencing of cell-culture isolates, which is based on combined results of up to six positive flasks per sample, with direct detection identifying fewer polioviruses (e.g. 8 or 9 versus 15 wild-type polioviruses and a lower prevalence of Sabin polioviruses).

**Figure 5.**
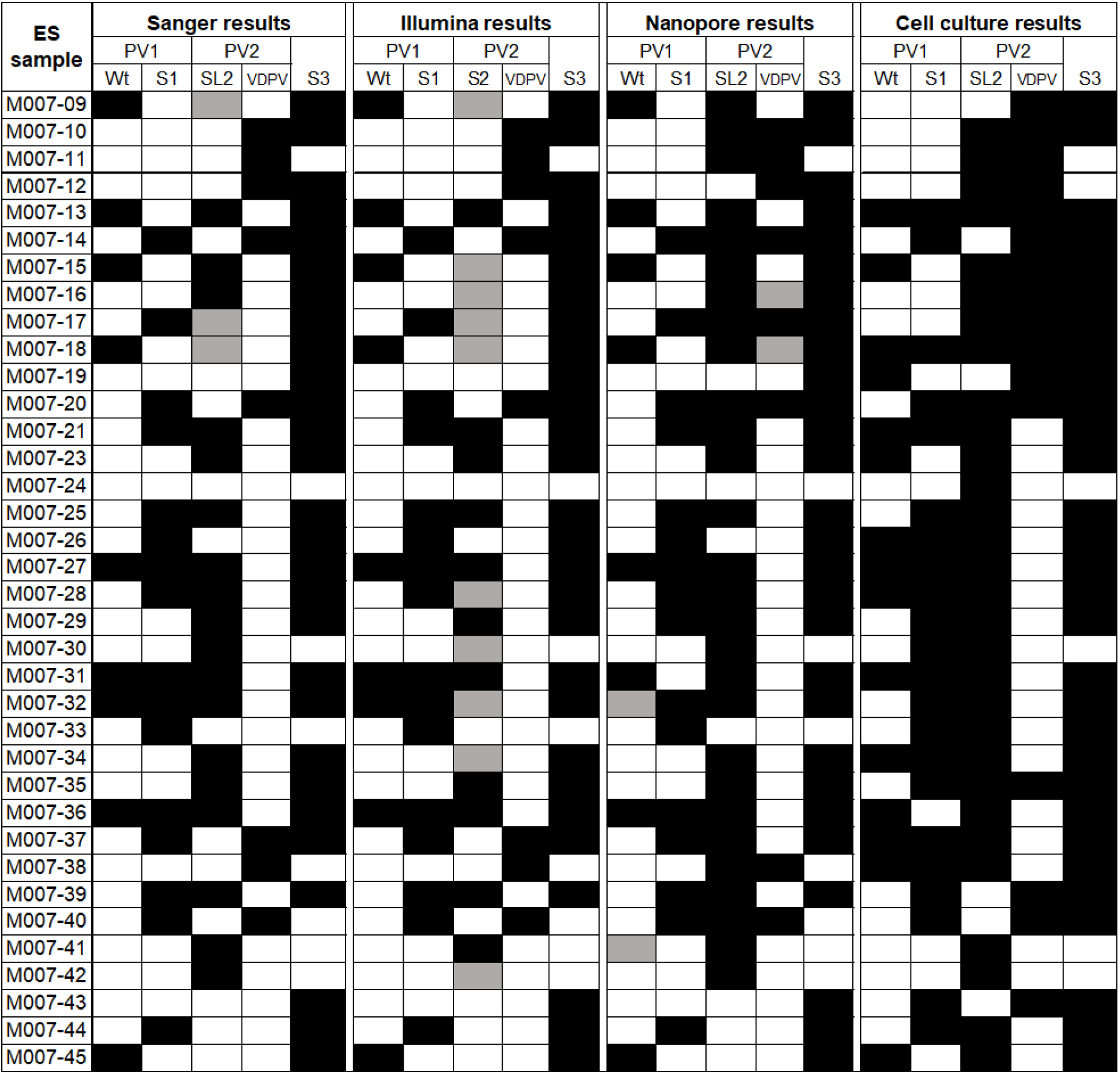
Polioviruses detected in Pakistan ES samples by cell culture or direct PCR followed by sequencing using three different methods. Detection of poliovirus in 36 environmental samples by nested PCR and nanopore sequencing compared with Illumina sequencing of the nested PCR product, serotype-specific PCR and Sanger sequencing, or Sanger sequencing of cell-culture isolates (up to 6 flasks per sample). A black cell indicates detection of the virus. Sabin-related detections have been clarified as either Sabin (S) or VDPVs where possible. For Sanger and Illumina results, grey indicates the detection of Sabin poliovirus and the possibility of a VDPV based on multiple peaks and SNP analysis respectively. For nanopore results, grey indicates detection of the virus but at a low read threshold.

Wild-type poliovirus nanopore consensus sequences were nearly identical to those obtained from the same sample aliquot by serotype-specific VP1 PCR and Sanger or Illumina sequencing (median identity to Sanger sequences of 100 % (range: 99.9-100.0%; n = 8)). These VP1 sequences were also very closely related to those obtained from corresponding cell-culture flasks using Sanger with a median identity of 100.0% (range: 98.7-100.0%); Figure 6 and Table S2). Similar results were obtained for Sabin 2 and VDPV2 sequences when compared to Sanger or Illumina sequencing results (median identity to Sanger sequences of 100.0% (range: 99.9 % to 100.0%, n = 26)).

**Figure 6.**
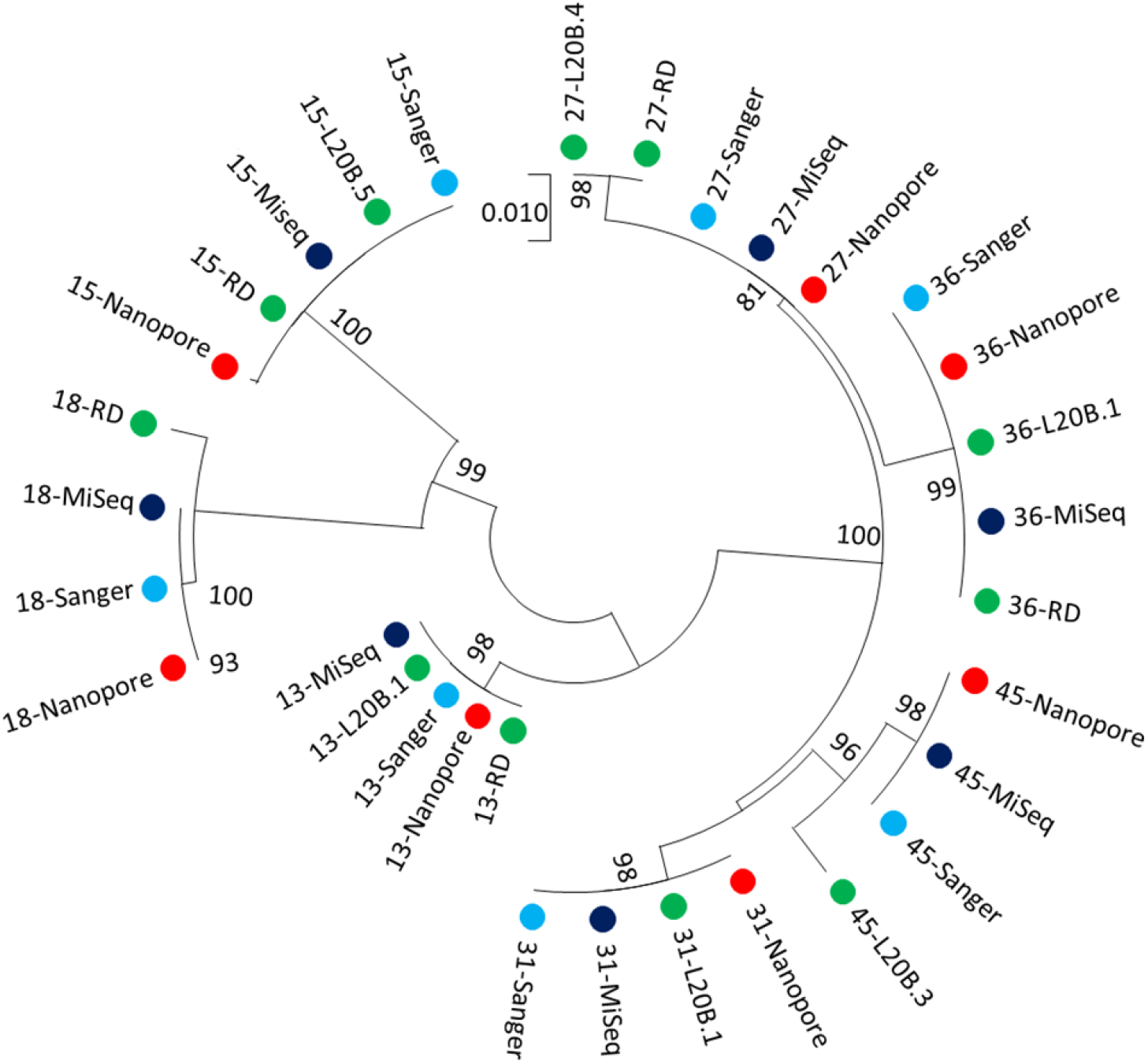
Relatedness of wild-type poliovirus 1 identified in ES samples in Pakistan using alternative methods. Consensus VP1 sequences derived by each sequencing platform (Sanger, MiSeq, nanopore) are shown in addition to the VP1 sequences of corresponding culture isolates (L20B, RD, labelled according to cell line used in isolation). No wild-type poliovirus 1 sequence was isolated by culture for sample M007-23, hence a isolate from a sample taken two months prior from the same ES site was used in the analysis (starred sequence).

### Real-time visualisation of demultiplexed sequencing results

The sequencing data generated through the nested VP1 protocol were analysed in real-time using RAMPART (Figure 7). This included live demultiplexing and mapping of each read to Sabin or wild-type polioviruses or NPEVs, allowing an immediate decision about when sufficient reads had been generated for each sample to permit ending of the sequencing run, preserving the MinION flow cell chemistry for further sequencing runs. At the end of each run, the custom bioinformatics pipeline created a report for each sample that describes the number of reads mapping to each reference and the corresponding consensus sequence.

**Figure 7.**
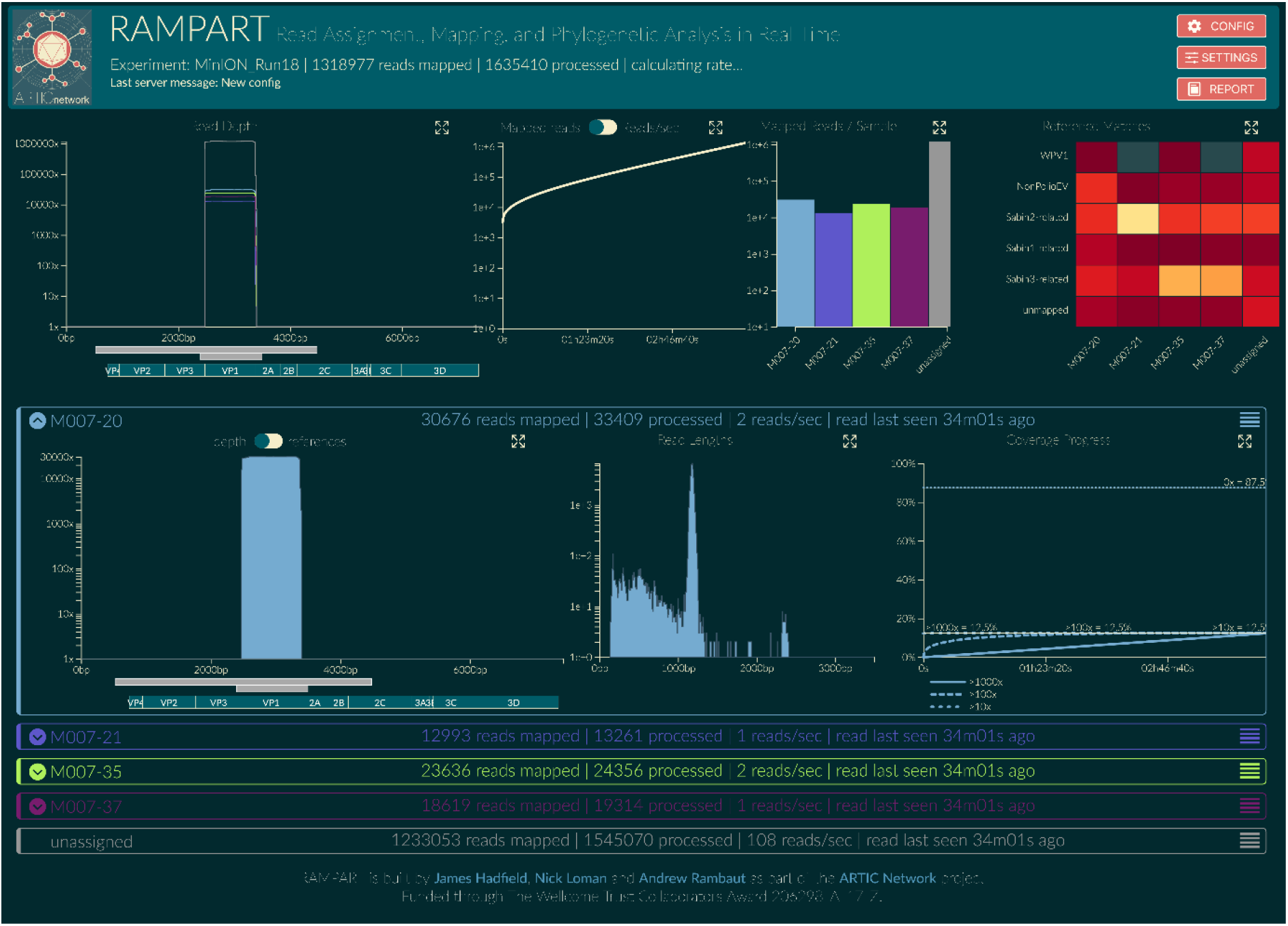
Illustration of RAMPART display for data from four ES samples. Top panel (left to right): 1) Summary of coverage across the genome for all samples, amplicons and genes of poliovirus. 2) Number of mapped reads plotted against time. 3) Number of mapped reads for each sample, unassigned reads shown in grey. 4) Heatmap showing proportion of reads for each sample that map to either wild-type poliovirus, Sabin-related poliovirus, NPEVs, or do not map (‘unmapped’). Lower panels: Detailed information can be displayed for each sample. The panel header indicates the sample that has been processed, the rate of data processing and the number of reads that have been mapped. Open panel (left to right): 1) Coverage across the genome and sample composition is displayed. 2) Read length distribution shown, with a peak at 1200 bases, the expected amplicon size. 3) Coverage of genome over time, which plateaus at 12.5% given that the amplicon spans only this proportion of the genome.

## Discussion

We present a culture-independent method for the rapid detection and sequencing of poliovirus from stool and ES samples. Using barcoded primers in a nested PCR, we were able to proceed from sample to sequence in three days, with real-time identification of polioviruses in each sample during nanopore (MinION) sequencing using RAMPART (15). The overall sensitivity of this method to detect wild-type or vaccine-derived polioviruses in Pakistan was 90% compared with culture, ITD and Sanger sequencing. Consensus sequences were near identical to those generated by Sanger sequencing of cell-culture isolates for the same samples, indicating that accurate VP1 sequences can be generated by nanopore sequencing. Further optimisation that we plan for this method has the potential to further increase sensitivity, such as testing duplicate extractions from a single stool sample and use of an RNA extraction control (e.g. bacteriophage Qβ).

Immediate identification of the poliovirus sequence allows downstream phylogenetic analysis and interpretation of epidemiological significance (30, 31). Faster generation of these results will allow earlier vaccination responses and containment of virus spread, which can have a substantially greater impact on poliomyelitis cases during an outbreak (32). Immediate availability of the poliovirus sequence allows potential sample contamination to be assessed and prevented - a challenge faced by direct detection methods relying on quantitative PCR, which requires subsequent additional PCR and sequencing reactions.

The high sensitivity of nested entire capsid/VP1 PCR for poliovirus in stool samples has been previously reported (100% sensitivity for 84 stool extracts (3)). Combining this approach with barcoded VP1 primers and nanopore sequencing allows sequencing of poliovirus mixtures and avoids the problem of non-specific amplification of NPEVs by the Q8/Y7 primers reported in (3) since any such amplicons are correctly classified as NPEVs by nanopore (unlike Sanger) sequencing.

Nanopore sequencing also allows mixtures of polioviruses and of polioviruses with other enteroviruses in stool and ES samples to be identified. In artificial mixtures, the distribution of sequencing reads was found to match the known composition of the mixture. Currently, identification of poliovirus mixtures requires laborious testing using multiple culture flasks, plaque isolation or antibody blocking of specific serotypes (33). Nanopore sequencing of ES samples gave results that were consistent with current culture-based methods but detected fewer polioviruses overall. This may reflect the much smaller aliquot of sewage concentrate (~200 μl) undergoing RNA extraction compared with that inoculated in cell-culture flasks (6 flasks with 0.5ml each). Indeed, testing the same RNA extract with RT-PCR using serotype-specific primers gave almost identical results and sequences to the nested PCR and nanopore sequencing, indicating that viruses within that aliquot were accurately and sensitively identified. Furthermore, the current cell-culture algorithm typically finds discordant results among flasks (34), in part due to heterogenous distribution of viruses within samples. This suggests that testing multiple aliquots or achieving greater sewage concentration would increase the sensitivity of direct molecular detection methods used to test ES samples.

Nanopore sequencing using MinION has a number of advantages over alternative next-generation sequencing approaches, which can also be applied to both culture supernatants and PCR products. These include lower initial investment and running costs, portability and ease of use. This has facilitated rapid training and implementation in laboratories using this technology for the first time. In addition, generation of full length reads allow shared mutations to be identified along the VP1 region, especially important for ES samples that may contain mixtures of polioviruses and where short reads often cannot distinguish multiple Sabin-related viruses.

The nested PCR and nanopore sequencing protocol can be scaled up to allow for the processing of multiple samples simultaneously, with groups of up to 96 samples being run together on a single MinION flow cell to provide the data included in this manuscript. This level of multiplexing still produced >15,000 reads per sample after just four hours of sequencing (using an R9.4.1 flow cell). Currently, the MinION flow cell chemistry can accommodate 48 hours of sequencing and therefore a single flow cell could in principle sequence >500 samples. In practice, we typically sequenced ~200 samples with a tail-off due to pore degradation.

The nested PCR and nanopore sequencing protocol has a number of limitations. Firstly, individual sequence reads on the MinION are error prone (currently ~5% single molecule error rate for R9 MinION flowcell (35)). An accurate consensus can be generated from just 50 reads or more, but for samples containing mixtures of closely related polioviruses (i.e. Sabin and VDPV of the same serotype) an initial clustering step is required. Our current bioinformatics pipeline was able to achieve accurate identification of Sabin, vaccine-derived and wild-type polioviruses in ES samples, but further validation is required. Sequencing errors may also affect accurate determination of sample barcodes. However, by enforcing stringent matching criteria (double barcodes with >80% identity) we observe negligible misclassification of reads (0.002%).

Secondly, a concern with repeated usage of flow cells is carry-over from one run to the next. This can however be prevented using a DNAse wash out and alternating sets of barcoded primers that would allow any carry-over to be identified (36). Thirdly, our protocol generates only VP1 sequences, consistent with current methods. However, full genome sequences may be of interest, and amplicons for the nearly full length genome can be generated directly from the sample and sequenced on MinION using pan-poliovirus primers (37). This PCR is less sensitive than the nested pan-EV-VP1 protocol and in some cases may require culture-based or other method of enrichment (e.g. (38)). Finally, careful handling of samples and laboratory processing is required to ensure optimal sensitivity and avoid sample cross-contamination. Further work is required to develop laboratory quality control and quality assurance procedures including provision of standardised positive controls if this method is to be adopted more widely by the Global Polio Laboratory Network. We demonstrate however that the immediate availability of sequencing data allows identification of contaminants and their removal from datasets, thus minimising false positives compared to other direct detection methods. Further optimisation of the method may also be possible, for example through testing alternative VP1 primers that are specific to particular polioviruses (e.g. for improved detection of wild-type 1 or circulating serotype 2 VDPV in ES samples).

In conclusion, we have developed a nested PCR with time and cost-saving barcoded primers that allows multiplex sample sequencing on a portable MinION sequencer. Using this method, only quantification, sample pooling and the addition of adaptors is required prior to sequencing. This method was shown to be sensitive and generate accurate poliovirus consensus sequences from stool samples. Complex poliovirus mixtures were identified in environmental samples, although further development of bioinformatic methods for consistent identification of variants within ES samples is required. This method has the potential to replace the current cell-culture based testing for poliovirus and has a number of significant advantages over alternative direct molecular detection methods.

## Acknowledgements

We would like to thank the members of the ARTIC network for helpful discussions in development of this work. We thank James Hadfield for support and development of the RAMPART software package.

## Funding

This work was funded by grants from the Bill and Melinda Gates Foundation (OPP1171890, OPP1207299). The Artic Network is funded by the Wellcome Trust through Collaborative Award 206298/Z/17/Z. NCG acknowledges support from the Medical Research Council (UK) Centre for Global Infectious Disease Analysis and a Royal Society Wolfson laboratory refurbishment scheme grant. AR acknowledges support from the European Research Council (grant agreement no. 725422-ReservoirDOCS).

## Notes

### Competing Interest Statement

The authors have declared no competing interest.

