## Supplemental Material for "Rapid and sensitive direct detection and identification of poliovirus from stool and environmental surveillance samples using nanopore sequencing"

**Figure S1 – Positions of primers used in this paper**

Positions relative to Toyoda *et al*. (1984)

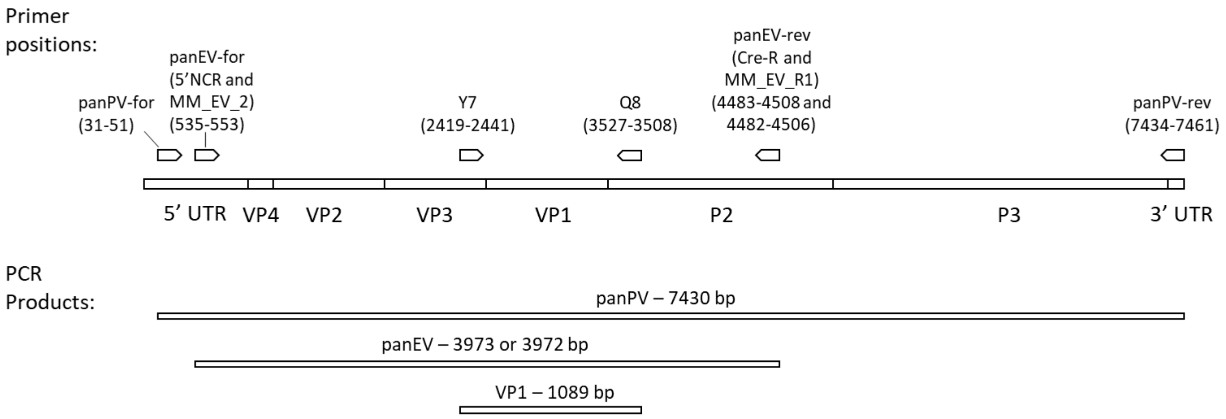

**Figure S2 – The accuracy of consensus sequences derived from varying sampling depths of a pure Sabin 1 sample sequenced using nanopore.**

Reads from a pure sample of Sabin 1 RNA derived from cell culture were sequenced via nanopore and the reads randomly sampled at varying depths. The resulting reads were mapped the VP1 region of Sabin 1 and polished to consensus using the module as outlined in the methods. For each number of reads drawn, 1,000 iterations were performed.

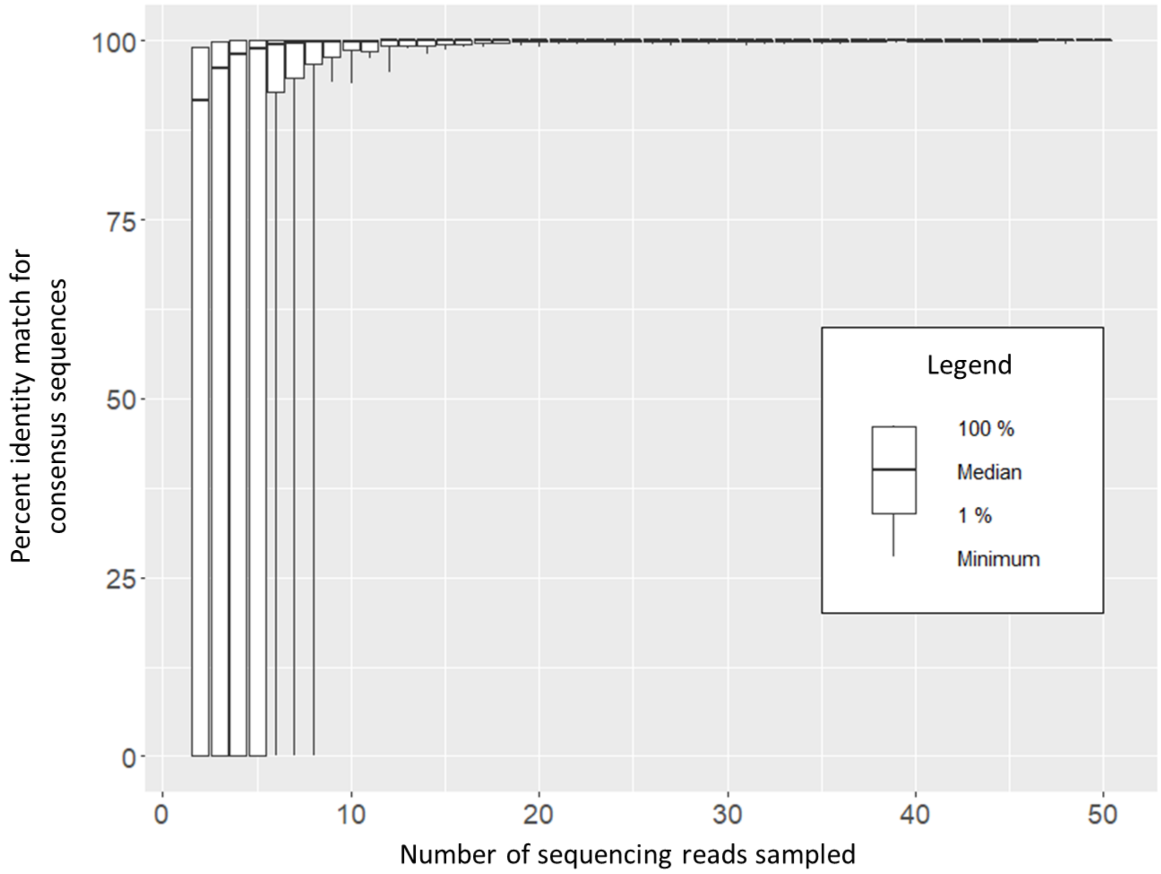


a) **The distribution of sequence identities for the consensus sequences compared to the Sabin 1 VP1 region.** Bars indicate the spread of the 99 % most accurate sequences with the central line indicating the median accuracy over the 1,000 iterations for that sampling depth. Whiskers indicate the spread of the remaining 1 % of reads i.e. the most inaccurate consensus sequences generated for each sequencing depth.

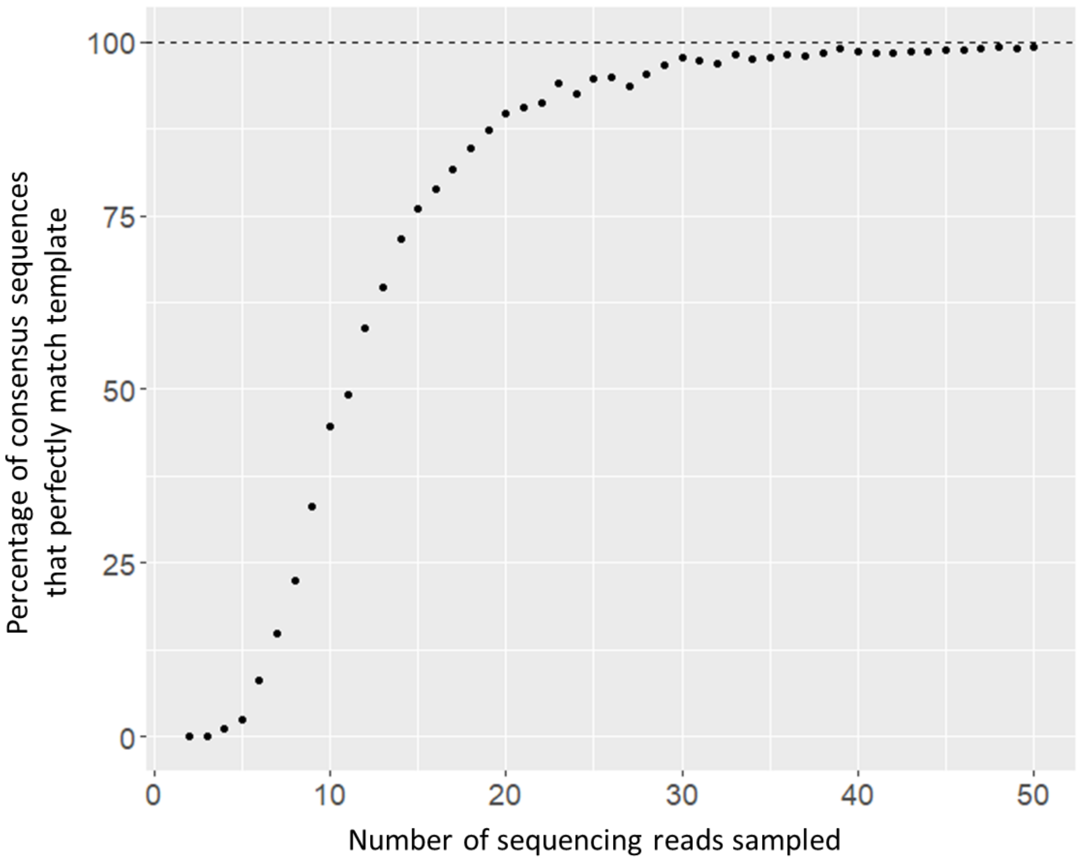


b) **The percentage of consensus reads generated at each sequencing depth that have 100 % identity to the Sabin 1 VP1 region.** For each sampling depth the percentage of consensus sequences over 1,000 iterations that perfectly match the Sabin 1 VP1 region is shown. 100 % would indicate that all consensus sequences at a chosen depth were a perfect match for Sabin 1; at a depth of 50 reads, 99.3 % of consensus sequences showed no mismatches when aligned to the Sabin 1 VP1 region.

**Table S1 - Summary of Runs used in this paper**

| Run | Purpose * | Run Duration (hours) | Number of Samples ** | Barcode use | Number of Active Pores | Total Mapped reads *** |
|---|---|---|---|---|---|---|
| **1** | PanEV mixtures (Figure 2) | 12 | 5 | 02-06 | 1340 | 133136 |
| **2** | PanEV Nanopore vs Illumina (Figure 3) | 14 | 12 | 01-12 | - | 628940 |
| **3** | ES samples run in a nested PCR with barcoded VP1 primers (Table2) | 2 | 12 | 26-36, 70 | 732 | 148252 |
| **4** | ES samples run in a nested PCR with VP1-BCA or VP1-barcoded primers | 3 | 38 | 43-80 | 1578 | 1115344 |
| **5** | ES samples run in a nested PCR with VP1 barcoded primers | 4 | 37 | 01-37 | 1438 | 424020 |
| **6** | Stool and ES samples run in a nested PCR with VP1 barcoded primers | 5 | 43 | 01-43 | 1033 | 438907 |
| **7** | ES samples run in a nested PCR with barcoded Y7R/Q8 primers | 5 | 36 | 13-48 | 947 | 338131 |
| **Pakistan1** | Stool and ES samples run in a nested PCR with VP1 barcoded primers | 2.5 | 48 | 01-48 | 1289 | 390142 |
| **Pakistan2** | Stool and ES samples run in a nested PCR with VP1 barcoded primers | 3 | 61 | 01-61 | 1426 | 265740 |
| **Pakistan3** | Stool samples run in a nested PCR with VP1 barcoded primers | 3 | 48 | 49-96 | 1126 | 428739 |
| **Pakistan4** | Stool and ES samples run in a nested PCR with VP1 barcoded primers | 4 | 56 | 41-96 | 969 | 271927 |
| **Pakistan5** | Stool and ES samples run in a nested PCR with VP1 barcoded primers | 2 | 83 | 01-83 | 1200 | 307011 |
| **India1** | Stool samples run in a nested PCR with Y7R/Q8 barcoded primers | 2 | 42 | 01-95 | 1140 | 195620 |
| **India2** | Stool samples run in a nested PCR with Y7R/Q8 barcoded primers | 2 | 50 | 01-94 | 1591 | 246137 |
| **India3** | Stool samples run in a nested PCR with Y7R/Q8 barcoded primers | 2 | 50 | 01-96 | 1656 | 779665 |

| | | | | | | |
|---|---|---|---|---|---|---|
| **India4** | Stool samples run in a nested PCR with Y7R/Q8 barcoded primers | 2 | 41 | 02-91 | 1656 | 688188 |
| **India5** | Stool samples run in a nested PCR with Y7R/Q8 barcoded primers | 2 | 54 | 01-90 | 1170 | 281381 |
| **India6** | Stool samples run in a nested PCR with Y7R/Q8 barcoded primers | 2 | 42 | 01-93 | 1070 | 207638 |
| **India7** | Repeat of select negative samples from India1-6 | 2 | 13 | 17-83 | 1055 | 6 |
| **India8** | Repeat of select negative samples from India1-6 | 2 | 3 | 17-81 | 1055 | 34 |

* Unless stated otherwise, VP1 PCR used the Y7/Q8 primer set

** This number includes Negative controls

*** Number of reads mapped to Poliovirus or NPEV from our database

**Table S2 - Sequence identities of consensus sequences compared between sequencing platforms**

| Sample | Nanopore Sequencing Reads | Nanopore versus | |
|---|---|---|---|
| | | Sanger | Illumina |
| M007-09 | 8212 | 100 | 100 |
| M007-10 | 919 | 100 | 100 |
| M007-11 | 12550 | 100 | 100 |
| M007-12 | 1641 | 100 | 100 |
| M007-13 | 2818 | 100 | 100 |
| M007-14 | 709 | 99.9 | 100 |
| M007-15 | 141 | 100 | 100 |
| M007-16 | 14644 | 100 | 100 |
| M007-17 | 3819 | 99.9 | 99.6 |
| M007-20 | 12565 | 100 | 100 |
| M007-23 | 10823 | 100 | 100 |
| M007-25 | 19926 | 100 | 100 |
| M007-27 | 5071 | 100 | 100 |
| M007-28 | 3528 | 100 | 100 |
| M007-29 | 32341 | 100 | 100 |
| M007-31 | 15597 | 100 | 100 |
| M007-32 | 24644 | 100 | 100 |
| M007-34 | 9388 | 100 | 100 |
| M007-35 | 7571 | 100 | 100 |
| M007-36 | 2866 | 100 | 100 |
| M007-37 | 7176 | 100 | 100 |
| M007-38 | 9079 | 100 | 100 |
| M007-39 | 1440 | 100 | 100 |
| M007-40 | 11451 | 100 | 100 |
| M007-41 | 14314 | 100 | 100 |
| M007-42 | 7719 | 100 | 100 |

a) Comparison of vaccine-derived poliovirus 2 consensus sequences. For each sample, the number of reads contributing to the nanopore consensus sequence is shown. Percentage identities between these sequences and consensus sequenced derived by direct sequencing using other sequencing platforms are shown.

| Sample | Nanopore Sequencing Reads | Nanopore versus | | | Sanger versus |
|---|---|---|---|---|---|
| | | Sanger | Illumina | Culture | Culture |
| M007-13 | 17531 | 100 | 100 | 100 | 100 |
| M007-15 | 3374 | 99.9 | 99.9 | 99.9 | 99.9 |
| M007-18 | 6464 | 100 | 100 | 99.2 | 99.2 |
| M007-27 | 14349 | 100 | 100 | 99.3 | 99.3 |
| M007-31 | 697 | 100 | 100 | 100 | 100 |
| *M007-32* | 25 | 100 | 100 | 100 | 100 |
| M007-36 | 1216 | 100 | 100 | 100 | 100 |
| *M007-45* | 3699 | 100 | 100 | 98.7 | 98.6 |

b) Comparison of wild-type poliovirus 1 consensus sequences. For each sample, the number of reads contributing to the nanopore consensus sequence is shown. Percentage identities between these sequences and consensus sequenced derived by direct sequencing using other sequencing platforms are shown, along with the sequence derived from Sanger sequencing of cultured isolates. The consensus of italicised Sanger sequences were truncated compared to other sequences; scoring was performed over aligned regions. Culture sequences were not available for sample 32; consensus sequences were compared to an isolate from the same ES site two months prior.

#### Appendix S1 – RNA extraction kit comparison experiments

1. **Method:**

Five RNA extraction kits were compared with extractions carried out as per the manufacturers' instructions. The kits tested were:

| **Kit name** | **Catalogue number** | **Input and output volumes** | **Notes** |
|---|---|---|---|
| Roche High Pure Viral RNA Kit (column-based method) | 11858882001 | 200 ul, RNA eluted in 50 ul | |
| QIAamp Viral RNA Mini Kit (column- based method) | 52904 | 140 ul, RNA eluted in 50ul | *As per protocol used two elution with 40 μL each time to maximise RNA recovery (80 ul total elution volume).* |
| MagMAX Viral RNA Isolation Kit (magnetic bead method). | AM1939 | 400ul, RNA eluted in 50 ul | |
| Zymo Viral RNA (column-based method) | R1035 | 800 ul, RNA eluted in 40 ul | *Zymo kit used according to CDC protocol[1], spun at 12000 x g in each step unless stated otherwise* |
| NucleoSpin RNA Virus F (column-based method). | 740958 | 1000 ul, RNA eluted in 50 ul | *NucleoSpin viral kit only used with 2016 samples* |

Eight samples extracted in total, all concentrates from sewage using the Two-Phase method:

From 2018: 18-031, 18-028, 18-019, 18-013

From 2016: 16-018, 16-019, 16-020, 16-021

RNA was stored at -80 °C after extraction and thawed on ice upon use. RNA was amplified in PanEV PCR reaction using both a three primer (Arita_rev, NIBSC R3, NIBSC F4) and a pair of two primer reactions (MM_EV_F2 and MM_EV_R1, Arita_for and Arita_rev) (sequence shown in Dataset S1).

#### 2. Results

For each kit and primer set, amplification was confirmed through gel electrophoresis. For the following figures, blue arrows indicate the height of the expected band.

##### 2.1. Roche kit – See Figures 1A and 1B

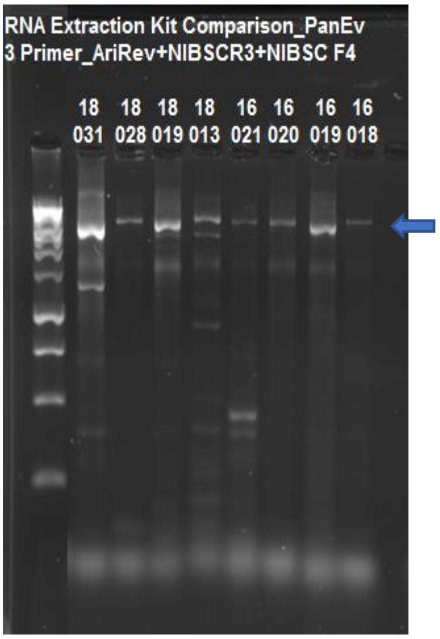


Figure 1A. Three Primer Reaction using Roche extracted RNA

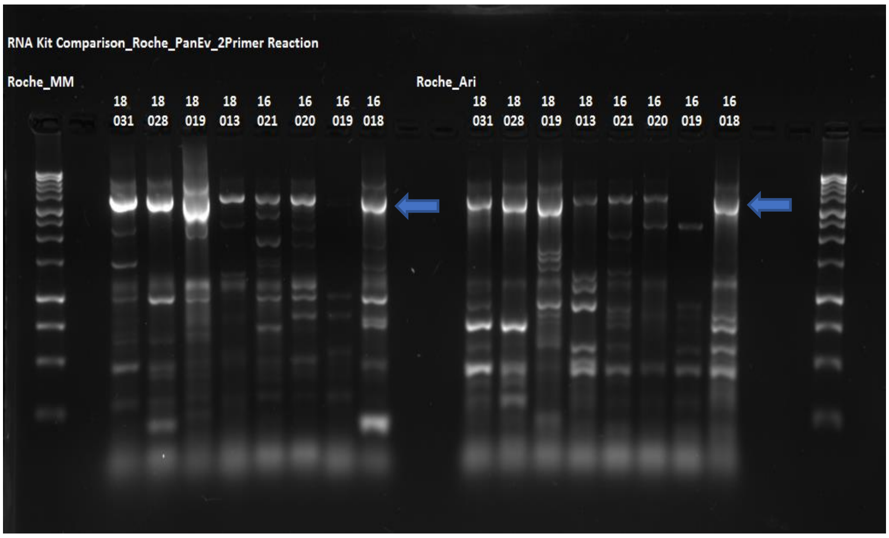


Figure 1B. Two Primer Reaction using Roche extracted RNA

2.2. MagMax kit – See Figures 2A and 2B

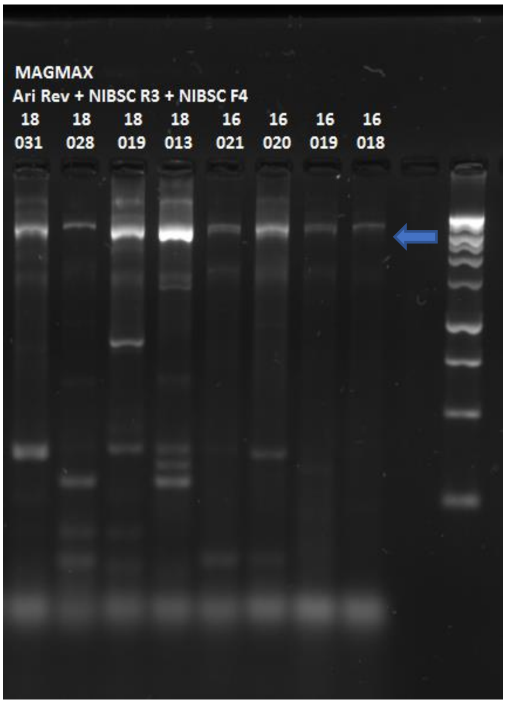


Figure 2A. Three Primer reaction using MagMax extracted RNA

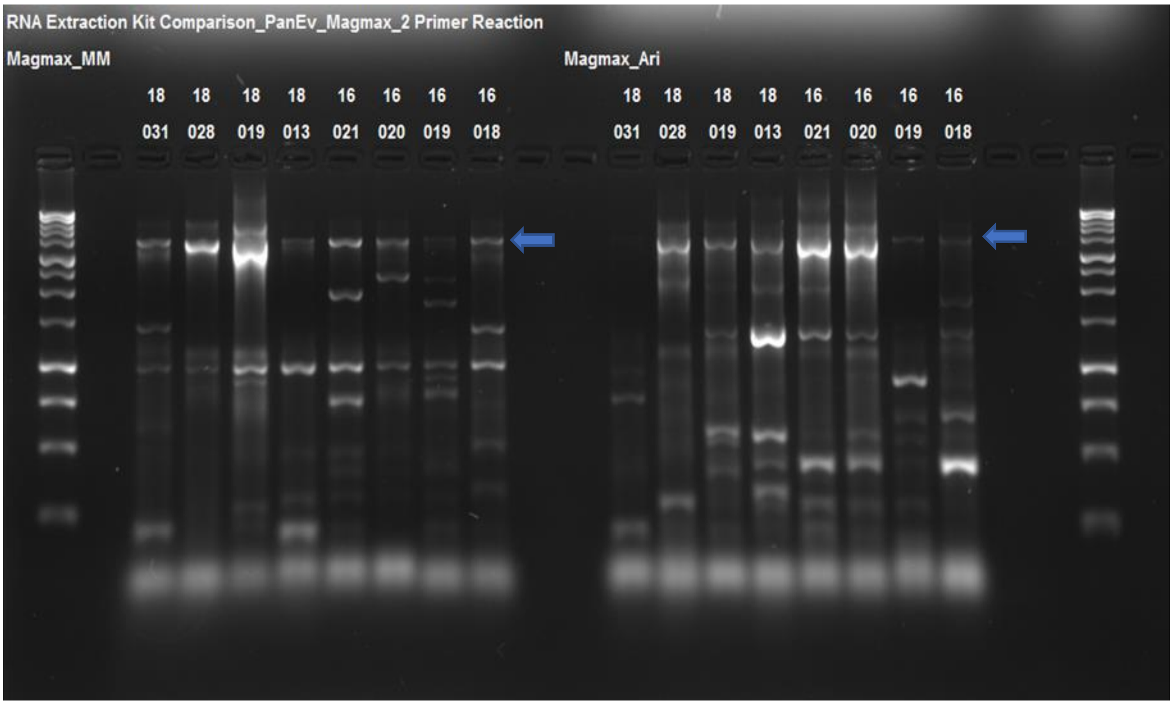


Figure 2B. Two Primer reaction using MagMax extracted RNA

2.3. Qiagen kit – See Figures 3A and 3B

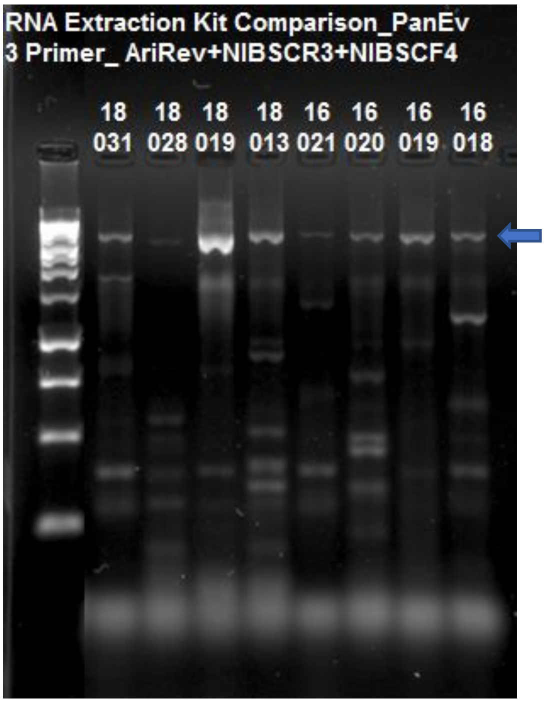


Figure 3A. Three Primer reaction using Qiagen extracted RNA

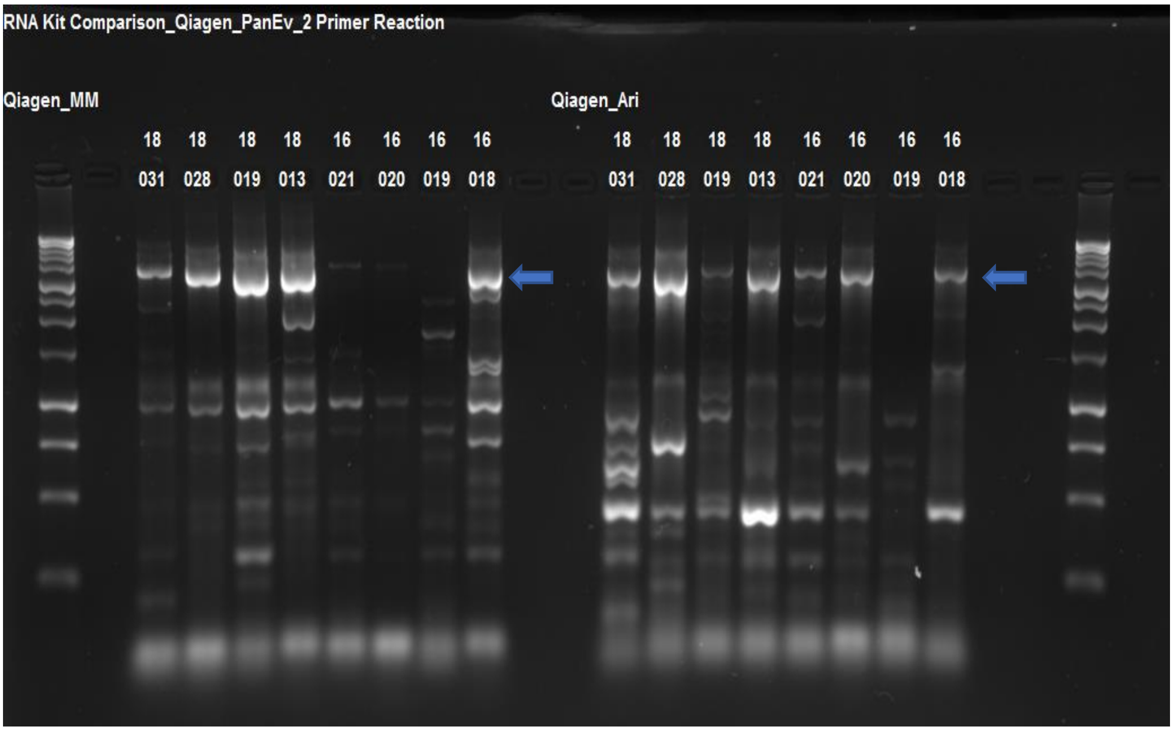


Figure 3B. Two Primer reaction using Qiagen extracted RNA

2.4. Zymo kit - See Figure 4

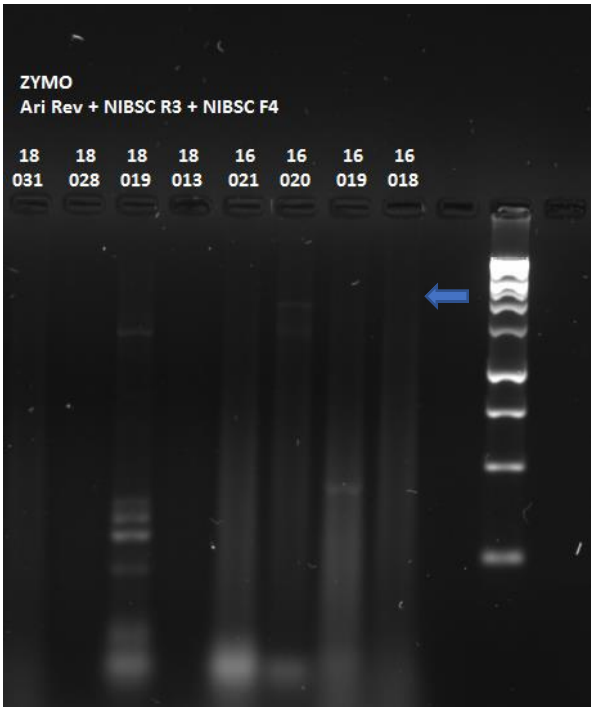


Figure 4. Three Primer reaction using Zymo extracted RNA

2.5. NucleoSpin kit – See Figure 5

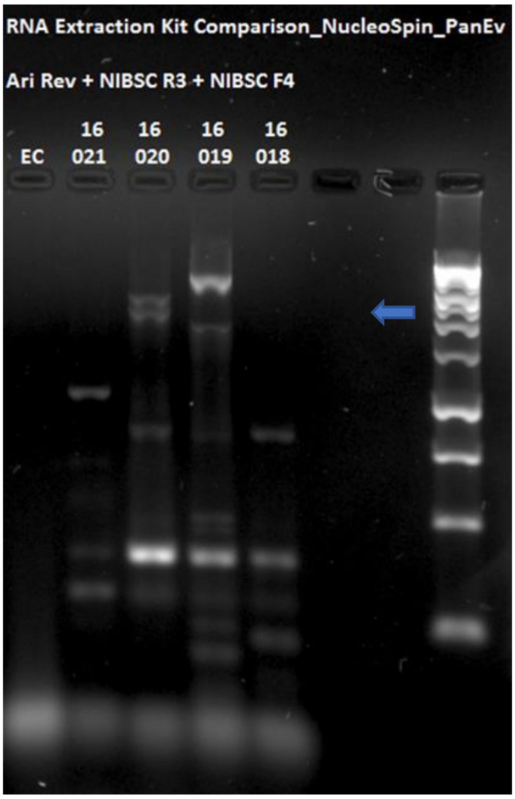


Figure 5. Three Primer reaction using NucleoSpin extracted RNA

#### 3. Conclusions

Long range PCR was observed to be possible with RNA extracted using column based or magnetic bead-based method. For the column-based methods, Roche and Qiagen each gave positive bands for 8 samples using the three primer reactions, whilst Roche performed better for the two primer reactions. Zymo and NucleoSpin gave poorer amplification. Amplification of MagMax extracted RNA gave similar results to that of the Roche kit.

**Appendix S2 - DNA Amplification and Sequencing Preparation Protocol**

---

**Version: 2.0**

**Last updated: 08/03/2020**

**Authors:**

The Vaccine Epidemiology Research Group, Imperial College London, UK

The Poliovirus Surveillance Group, National Institute for Biological Standards and Control, South Mimms, UK

#### Consumables

SuperScript™ III One-Step RT-PCR System (ThermoFisher, 12574018)

Blunt/TA Ligase Master Mix (NEB, M0367) *Only required for standard primers*

PCR Barcoding Kit (Oxford Nanopore, EXP-PBC001) or PCR Barcoding Expansion Pack 1-96 (Oxford Nanopore, EXPPBC096) *Not required for barcoded primers*

Flow Cell Priming Kit (Oxford Nanopore, EXP-FLP001)

Agencourt AMPure XP beads (Beckman Coulter)

Ligation Sequencing Kit 1D (Oxford Nanopore, SQK-LSK109)

LongAmp Taq 2X Master Mix (NEB, M0287) or DreamTaq 2x Master Mix (K1072)

NEBNext End repair / dA-tailing Module (NEB, E7546)

NEBNext FFPE DNA Repair Kit (NEB, M6630)

NEBNext Quick Ligation Module NEB, (E6056)

Freshly prepared 70% ethanol in nuclease-free water

Nuclease-free water

#### References

The section "Library Preparation for the ONT MinION" is based on the Oxford Nanopore Technologies 1D PCR barcoding amplicon/cDNA (SQK-LSK109).

### DNA amplification

**For Standard PCR- Complete steps 1 to 5 and step 13**

**For Nested VP1 PCR- Complete steps 1 to 13**

1) Prepare a Master mix using reaction volumes as detailed below, excluding forward primer and the RNA:

| | 1 Reaction (µL) | Reactions |
|---|---|---|
| 2x Master Mix | 12.5 | µL |
| SS III Platinum Taq mix | 0.5 | µL |
| Reverse Primer (10 µM) | 1 | µL |
| Nuclease Free Water | 5 | µL |
| Forward Primer (10 µM) | 1 | - |
| RNA | 5 | - |
| Total volume | 25 | |

2) Briefly vortex and centrifuge down. Add 19 µL of master mix to each PCR tube and 5 µL of eluted RNA.

3) Incubate at 50 °C for 30 minutes.

4) Add 1 µL of the forward primer to the tubes.

5) Amplify using the following cycling conditions:

| **CYCLE** | **STEP** | **TEMP (°C)** | **TIME** |
|---|---|---|---|
| 1 | Initial Denaturation | 94 | 2 minutes |
| 42 | Denaturation | 94 | 15 seconds |
| | Annealing | 55 | 30 seconds |
| | Extension | 68 | 4 minutes 30 seconds* |
| 1 | Final Extension | 68 | 5 minutes |
| - | Hold | 10 | - |

*Extension time for panEV amplification

6) Dilute 5 µL of PCR product in 95 µL of nuclease free water and vortex

7) Prepare a Master mix using reaction volumes as detailed below, excluding the forward primer and diluted PCR product (and reverse primer if barcoded):

| | 1 Reaction (µL) | Reactions |
|---|---|---|
| DreamTaq 2x master mix | 12.5 | µL |
| Water | 5.5 | µL |
| Forward primer (10 µM) | 1 | - |
| Reverse primer (10 µM) | 1 | µL |
| Diluted PCR product | 5 | - |
| Total volume | 25 | |

8) Briefly vortex and centrifuge down the master mix and aliquot 18 µL into each PCR tube.

9) Add 1 µL of applicable primers.

10) Add 5 µL of the diluted DNA.

11) Amplify using the following cycling conditions:

| CYCLE | STEP | TEMP (°C) | TIME |
|---|---|---|---|
| 1 | Initial Denaturation | 94 | 2 minutes |
| 35 | Denaturation | 94 | 15 seconds |
| | Annealing | 55 | 30 seconds |
| | Extension | 65 | 2 minutes |
| 1 | Final Extension | 65 | 10 minutes |
| - | Hold | 10 | - |

12) PCR confirmation- Check a representative set of samples to confirm that the PCR has been successful.

13) AMPure bead purification

- Prepare the AMPure XP beads for use; resuspend by vortexing.
- Add 15 µL (0.6 ratio) of resuspended AMPure XP beads to the reaction and mix by pipetting.
- Incubate on a rotator for 5 minutes at room temperature.
- Spin down the sample and pellet on a magnet. Keep the tube on the magnet, and pipette off the supernatant.
- Keep on magnet, wash beads with 100 µL of freshly prepared 70% ethanol without disturbing the pellet. Remove the 70% ethanol using a pipette and discard. Repeat.
- Spin down and place the tube back on the magnet. Pipette off any residual 70% ethanol. Briefly allow to dry.
- Remove the tube from the magnetic rack and resuspend pellet in 25 µL nuclease-free water. Incubate for 2 minutes at RT.
- Pellet beads on magnet until the eluate is clear and colourless.
- Remove and retain 23 µL of eluate in a clean 1.5 mL Eppendorf DNA LoBind tube.
- Quantify the eluted DNA.

### Library Preparation for the ONT MinION

**For Standard Primers- Complete all steps**

**For BCA primers- Complete steps 4 to 10**

**For Barcoded primers- Complete steps 7 to 10**

#### 1) Standardise DNA

- Transfer 1 µg of DNA into a clean PCR tube.
- Adjust the volume to 23 µL with nuclease-free water.

#### 2) End-prep & dA-tailing

- Prepare the following reaction mix:

  23 µL <1 µg DNA

  3.5 µL Ultra II End-prep reaction buffer

  1.5 µL Ultra II End-prep enzyme mix

  2.5 µL Nuclease-free water

- Mix gently by flicking, and spin down.
- Incubate for 5 minutes at 20 °C and 5 minutes at 65 °C.
- Prepare the AMPure XP beads for use; resuspend by vortexing.
- Add 30 µL of resuspended AMPure XP beads to the end-prep reaction and mix by pipetting.
- Incubate on a rotator for 5 minutes at room temperature.
- Spin down the sample and pellet on a magnet. Pipette off the supernatant.
- Wash beads with 100 µL of freshly prepared 70% ethanol without disturbing the pellet.
- Remove the 70% ethanol using a pipette and discard. Repeat.
- Spin down and place the tube back on the magnet. Pipette off any residual 70% ethanol. Briefly allow to dry.
- Remove the tube from the magnetic rack and resuspend pellet in 20 µL Nuclease-free water.
- Incubate for 2 minutes at room temperature.
- Pellet beads on magnet until the eluate is clear and colourless.
- Remove and retain 16 µL of eluate into a clean PCR tube.
- Quantify 1 µL of end-prepped DNA using a Qubit fluorometer - recovery aim > 700 ng.

#### 3) Ligation of Barcode Adapter

- Add the reagents in the order given below:

  15 µL End prep DNA

  10 µL Barcode Adapter

  25 µL Blunt/TA Ligase Master Mix

- Mix gently by flicking the tube, and spin down.
- Incubate the reaction for 10 minutes at room temperature.
- Prepare the AMPure XP beads for use; resuspend by vortexing.
- Add 20 µL of resuspended AMPure XP beads to the reaction and mix by pipetting.
- Incubate on a rotator for 5 minutes at room temperature.
- Spin down the sample and pellet on a magnet. Pipette off the supernatant.
- Keep on magnet, wash beads with 100 µL of freshly prepared 70% ethanol without disturbing the pellet. Remove the 70% ethanol using a pipette and discard. Repeat.
- Spin down and place the tube back on the magnet. Pipette off any residual 70% ethanol. Briefly allow to dry.
- Remove the tube from the magnetic rack and resuspend pellet in 25 µL nuclease-free water.
- Incubate for 2 minutes at room temperature.
- Pellet beads on magnet until the eluate is clear and colourless.
- Remove and retain 15 µL of eluate in a clean tube.
- Quantify 1 µL of DNA.

#### 4) Standardise DNA

- Transfer 100 fmol of DNA into a clean PCR tube.
- Adjust the volume to 24 µL with nuclease-free water.

#### 5) Barcoding PCR

- Set up a barcoding PCR reaction as follows for each sample:

  1 µL PCR Barcode

  24 µL 100 fmol PCR Product

  25 µL LongAmp Taq 2x master mix

- Mix gently by flicking the tube, and spin down.
- Amplify using the following cycling conditions:

| CYCLE | STEP | TEMP (°C) | TIME |
|---|---|---|---|
| 1 | Initial Denaturation | 95 | 3 minutes |
| 12 | Denaturation | 95 | 15 seconds |
| | Annealing | 62 | 15 seconds |
| | Extension | 65 | 2 minutes* |
| 1 | Final Extension | 65 | 2 minutes |
| - | Hold | 4 | - |

*Extension time for VP1 amplification

#### 6) Fragment Purification

- Prepare the AMPure XP beads for use; resuspend by vortexing.

- Add 40 µL (0.8 ratio) of resuspended AMPure XP beads to wells and mix by pitpetting.
- Incubate on a rotator for 5 minutes at room temperature.
- Spin down the sample and pellet on a magnet. Keep the tube on the magnet, and pipette off the supernatant.
- Keep on magnet, wash beads with 100 µL of 70% ethanol without disturbing the pellet. Remove the 70% ethanol using a pipette and discard. Repeat.
- Spin down and place the tube back on the magnet. Pipette off any residual 70% ethanol. Briefly allow to dry.
- Remove the tube from the magnetic rack and resuspend pellet in 25 µL nuclease-free water. Incubate for 2 minutes at room temperature.
- Pellet beads on magnet until the eluate is clear and colourless.
- Remove and retain 23 µL of eluate in a clean PCR plate.

#### 7) Sample Pooling

- Prepare 1 µg of pooled barcoded DNA in 47 µl Nuclease-free water.

Optional- If pool volume >47 µL concentrate with the following step:

- Prepare the AMPure XP beads for use; resuspend by vortexing.
- Add the required amount of resuspended AMPure XP beads to the DNA pool and mix by pipetting.
- Incubate on a rotator for 5 minutes at room temperature.
- Spin down the sample and pellet on a magnet. Keep the tube on the magnet, and pipette off the supernatant.
- Keep on magnet, wash beads with 100 µL of freshly prepared 70% ethanol without disturbing the pellet. Remove the 70% ethanol using a pipette and discard. Repeat.
- Spin down and place the tube back on the magnet. Pipette off any residual 70% ethanol. Briefly allow to dry.
- Remove the tube from the magnetic rack and resuspend pellet in 50 µL nuclease-free water. Incubate for 2 minutes at room temperature.
- Pellet beads on magnet until the eluate is clear and colourless.
- Remove and retain 47 µL of the eluate in a clean 1.5 ml Eppendorf DNA LoBind tube.

#### 8) End-prep and dA-tailing

- Add the following reagents in a 0.2 mL PCR tube.

  47 µL 1 µg DNA

  3.5 µL NEBNext FFPE DNA Repair Buffer

  2 µL NEBNext FFPE DNA Repair Mix

  3.5 µL Ultra II End-prep reaction buffer

  3 µL Ultra II End-prep enzyme mix

  1 µL DNA CS

- Mix gently by flicking the tube, and spin down.
- Incubate for 10 minutes at 20 °C and 10 minutes at 65 °C.
- Place on ice for 30 seconds.
- Transfer sample to a 1.5 mL Eppendorf DNA LoBind tube.
- Prepare the AMPure XP beads for use; resuspend by vortexing.
- Add 60 µL of resuspended AMPure XP beads to the end-prep reaction and mix by pipetting.
- Incubate on a rotator for 5 minutes at room temperature.
- Spin down the sample and pellet on a magnet. Keep the tube on the magnet, and pipette off the supernatant.
- Keep on magnet, wash beads with 200 µL of freshly prepared 70% ethanol without disturbing the pellet. Remove the 70% ethanol using a pipette and discard. Repeat.
- Spin down and place the tube back on the magnet. Pipette off any residual 70% ethanol. Briefly allow to dry.
- Remove the tube from the magnetic rack and resuspend pellet in 61 µL Nuclease-free water. Incubate for 2 minutes at room temperature.
- Pellet beads on magnet until the eluate is clear and colourless.
- Remove and retain 60 µL of eluate into a clean 1.5 mL Eppendorf DNA LoBind tube.

**9) Adaptor ligation**

- Thaw and prepare the kit reagents as follows:

  Spin down and thaw Adapter Mix (AMX) on ice

  Spin down and thaw T4 Ligase from NEBNext Quick Ligation Module (E6056) on ice
  Thaw Ligation Buffer (LNB) at room temperature, spin down, mix by pipetting. Place on ice.
  Thaw Elution Buffer (EB) at room temperature, mix by vortexing, spin down. Place on ice.
  Thaw one tube of L Fragment Buffer (LFB) for panEV or S Fragment Buffer (SFB) for VP1 at room temperature, mix by vortexing, spin down and place on ice

- Prepare the following reaction mix in a 1.5 mL Eppendorf DNA LoBind tube:

  60 µL DNA

  25 µL Ligation Buffer (LNB)

  10 µL NEBNext Quick T4 DNA Ligase

  5 µL Adapter Mix (AMX)

- Mix gently by flicking the tube, and spin down.
- Incubate the reaction for 10 minutes at room temperature.

**10) AMPure XP bead binding**

- Prepare the AMPure XP beads for use; resuspend by vortexing.
- Add 40 µL of resuspended AMPure XP beads to the adapter ligation reaction from the previous step and mix by pipetting.
- Incubate on a rotator for 5 minutes at room temperature.
- Place on magnetic rack, allow beads to pellet and pipette off supernatant.

- Add 250 µL of the LFB/SFB to the beads. Close the tube lid and resuspend the beads by flicking the tube. Return the tube to the magnetic rack, allow beads to pellet and pipette off the supernatant. Repeat.
- Spin down the tube and place back on the magnet. Pipette off residual supernatant and briefly air dry.
- Remove the tube from the magnetic rack and resuspend pellet in 15 µl Elution Buffer. Incubate for 10 minutes at room temperature.
- Pellet beads on magnet until the eluate is clear and colourless.
- Remove and retain the eluate which contains the DNA library in a clean 1.5 mL Eppendorf DNA LoBind tube
- Dispose of the pelleted beads

The prepared library is used for loading into the flow cell. Store the library on ice until ready to load.

**Appendix S3 – Comparison of the effect of VP1 primer modification by addition the barcode adapter or barcode sequences.**

A dilution series of PV1 RNA was set up starting from a $10^{-1}$ dilution of the RNA concentrate down to a $10^{-7}$ dilution. RT-PCR was carried out as previously described (Appendix S2), and subsequent VP1 PCR nest was carried out with the Y7/Q8 primer set, the primers with the barcode adapter (VP1-BCA) or with ONT barcode sequence (VP1-BC70) appended to the original primer sequence (all primer sequences in Table S1). PCR products, including those from the RT-PCR, were visualised on a 1% agarose gel stained with SYBRsafe (Invitrogen) and run at 120V for 1hour with a 1kb plus DNA ladder (NEB, Figure 1).

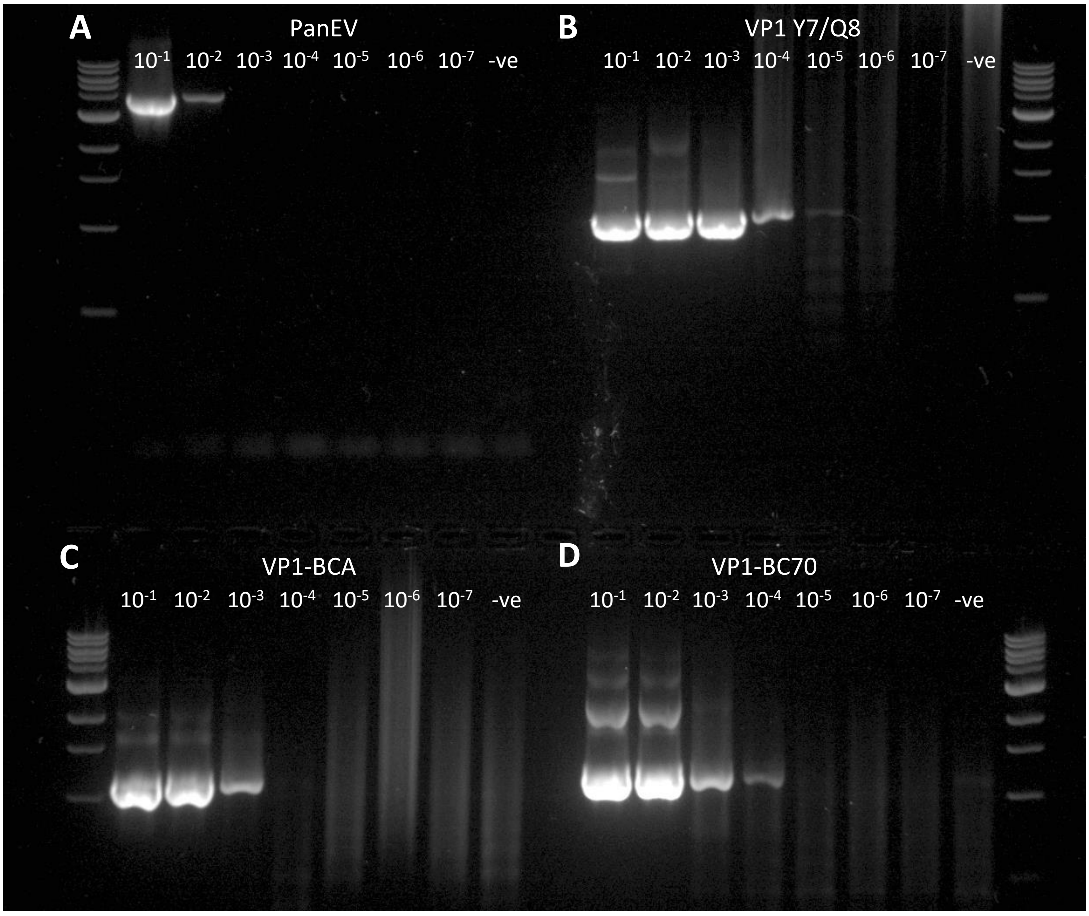


*Figure 1: Visualisation of products from: a) PanEV RT-PCR, b) plain Y7/Q8 PCR, c) Y7/Q8 primers with the barcoding adapter, and d) Y7/Q8 primers with ONT barcode 70*

Remaining PCR products up to the $10^{-6}$ dilution factor from the plain VP1 Y7/Q8 reaction and the VP1-BCA reaction were processed further to assess how much product was lost during the additional steps required to get the products up to the same point as the products in the VP1-BC70 reaction (method in Appendix S2). The products after the barcoding PCR were visualised as before (Figure 2).

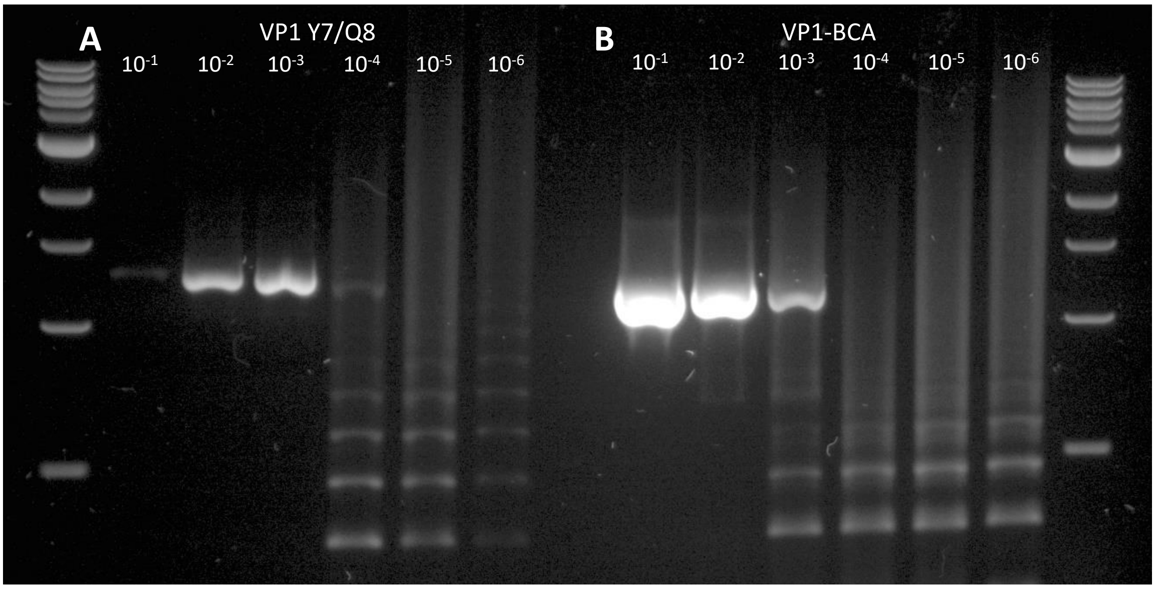


*Figure 2: Visualisation of products from Y7/Q8 PCR products (a) and VP1-BCA products (b) continued to the barcoding PCR stage of library preparation for sequencing on the MinION.*

After the barcoding PCR, Y7/Q8 products were seen up to the $10^{-4}$ dilution of original input RNA; for the VP1-BCA products, a band was only seen up to the $10^{-3}$ dilution (Figure 2). In the original PCR nest, the barcoded primers gave a visible product up to the $10^{-4}$ dilution (Figure 1), with a slightly brighter band being produced in comparison to the Y7/Q8 products after the barcoding PCR. This implies that there is little difference in sensitivity of the Y7/Q8 primers – which would then have to go through the full library preparation process – and of the barcoded primers which offer a much quicker option for library preparation.

**Appendix S4 – Additional samples tested with the nested VP1 protocol and nanopore sequencing in Vellore, India.**

Testing of stored stool samples in India, which had detectable (Ct<40) serotype 1 or 3 Sabin poliovirus by serotype-specific qRT-PCR in 2013, yielded >50 reads of the correct virus in 57% (58/102) and 79% (91/115) of samples respectively when using nested PCR with barcoded Y7R/Q8 VP1 primers and nanopore sequencing. Detection was strongly dependent on viral copy number measured by the original qRT-PCR (Wilcoxon rank sum test, $p<0.001$). Among samples with at least 100 copies of the target sequence at the time of original testing, 87% (40/46) and 93% (69/74) were positive for serotypes 1 and 3 by nanopore sequencing (>50 reads). After accounting for variability among samples in viral copy number using multivariable logistic regression, the probability of detecting serotype 3 poliovirus by nanopore sequencing was somewhat higher compared with serotype 1 (odds ratio 1.12; 95% confidence interval 1.01 – 1.25). Culture done in 2013 was positive for 87% (89/102) and 89% (102/115) of these samples respectively. None of the 49 samples negative by qRT-PCR and culture were positive for poliovirus by nested PCR and nanopore sequencing.
